# The consequences of array size for human centromere performance in mitosis

**DOI:** 10.64898/2026.06.15.732503

**Authors:** Ayantika Sen Gupta, Siddharth Shivanandan, Mark Mattingly, Jay R. Unruh, Sean A. McKinney, Arundhathy Prasanthy Jithesh, Simona Giunta, Jennifer L. Gerton

**Affiliations:** Stowers Institute for Medical Research, Kansas City, Missouri, USA; Giunta Laboratory of Genome Evolution, Department of Biology and Biotechnologies Charles Darwin, University of Rome “Sapienza”, Piazzale Aldo Moro 5, 00185 Rome, Italy; Indian Institute of Science Education and Research, Pune, India

## Abstract

Long-read sequencing and assembly of human genomes have revealed extreme variation in centromeric alpha-satellite array sizes across chromosomes and individuals. However, assessing the impact of centromeric array size on centromere function remains challenging. In this study, using quantitative FISH and microscopy assays, we established a framework for the systematic functional evaluation of naturally occurring variations in centromeric array size during mitosis. By comparing 8 pairs of homologous chromosomes exhibiting 1.24 – 4.6 –fold variation in centromeric array size, we uncovered size-based differences in function. We found that between a pair of homologous chromosomes, the chromosome harboring the smaller centromeric array is more prone to chromosome missegregation. Centromere size-based differences in chromosome missegregation rates cannot be explained by differential enrichment of molecular factors like centromeric histone CENP-A, CENP-B, or most kinetochore proteins. However, the smaller centromere out of a homologous pair consistently exhibits increased cohesion fatigue, suggesting the involvement of the cohesin complex in size-based differences in centromere cohesion. Consistent with our hypothesis, complete deprotection of the cohesin complex by depletion of shugoshin-1, removes the size-based bias in centromere cohesion. These results suggest that while CENP-A–enriched core centromeres are essential for kinetochore assembly, variation in overall centromeric array size has a significant impact on centromere performance and the fidelity of chromosome segregation during mitosis.

## Introduction

Centromeres are rapidly evolving, epigenetically defined regions on chromosomes that are required for accurate sister chromatid segregation between daughter cells during cell division^1–3^. Incorporation of the centromeric histone H3 variant CENP-A, assembly of the large protein complex known as the kinetochore, and attachment of the kinetochore to microtubules are all essential processes for accurate sister chromatid segregation during mitosis^4^. Moreover, sister chromatids must remain cohesed until biorientation is achieved and sister chromatids can be accurately segregated. Cohesion at centromeres depends on the cohesin complex and DNA catenations^5–7^. Decades of research has described many of the protein components and processes required for accurate sister chromatid segregation, but much less is known about how centromeric DNA itself contributes to chromosome transmission.

In humans, native centromeres are composed of highly repetitive α-satellite DNA consisting of 171-bp monomers^8^. α-satellite monomers are arranged into higher-order repeats or HORs, a stereotypical series of sequence distinct monomers that are chromosome-specific^9,10^. HORs are 97-100% identical at the sequence level on homologous chromosomes, but only 70% identical between chromosomes^11–13^. Recent human genome assemblies have demonstrated large-scale polymorphisms in the size of α-satellite centromeric DNA arrays between chromosomes, homologs and individuals^14–16^. Furthermore, repression of meiotic recombination in and around centromeres means that centromeres are inherited intact through the germline in the human population, termed centromere haplotypes^17–19^. A previous study used cytogenetics to demonstrate that centromere 17 epialleles composed of more α-satellite DNA repeats outcompete shorter epialleles for enrichment of the centromeric histone CENP-A^20^. However, it is unclear how other centromere functions relate to centromeric array size.

Chromosomes exhibit non-random rates of missegregation and aneuploidy in different cell types and contexts^21–25^. However, a limitation of these prior works is that they analyzed aneuploidy frequency in relation to centromere size but by averaging homologs and comparing structurally distinct chromosomes. Thus, the impact of centromeric array size on chromosome segregation has not been examined in isolation. Due to the highly repetitive and heterochromatic nature of centromeres, genetic manipulation of these sites to create size variants under isogenic conditions remains challenging. Here we bypass this challenge by comparing centromere function between two homologous chromosomes with distinct centromeric array sizes within a single cell. We directly visualized centromere size using quantitative fluorescence in-situ hybridization (qFISH). We benchmarked our method using the newly available telomere-to-telomere assembly of the RPE-1 genome^26,27^. We identified chromosome-specific centromeric array size variants across cell lines and compared their function during mitosis. The homolog bearing the shorter centromeric array missegregates more frequently when the spindle checkpoint is compromised. Furthermore, shorter centromeric arrays are less cohesed compared to longer centromeric arrays but perturbation of centromeric cohesin equalizes their performance. Overall, our results indicate that centromeric array size can be a genetic determinant of the mitotic function of centromeres and contribute to aneuploidy.

## Results

### Centromeric array size can be distinguished by FISH in human cell lines

To understand the impact of array size variation on centromere performance, we first needed to establish a quantitative FISH approach that could reliably identify centromeres on homologous chromosomes that differed in their sizes. Most human chromosomes, with the exception of 1, 5, 13, 14, 19, 21 and 22, can be uniquely identified by their centromere-specific sequences, detected by fluorescence *in-situ* hybridization (FISH) probes. DNA FISH has been used to discriminate between homologous sites on chromosomes that vary in size, including telomeres and the chromosome 15 pericentromeric HSat III array^28,29^. We used this quantitative FISH (qFISH) approach to identify size variation between homologous chromosomes, focusing on the active alpha-satellite HOR array that forms the core centromere and pericentromeric domains. The telomere-to-telomere assembled diploid RPE-1 reference genome revealed many homologous pairs with variation in centromere array size (Figure 1A). Based on the RPE-1 genome assembly, we found chromosomes like chromosome 1, with no variation in array size between homologs, and chromosomes like chromosome 21, with nearly 4.6-fold variation in centromeric array size between homologs (Figure 1B).

**Figure 1.**
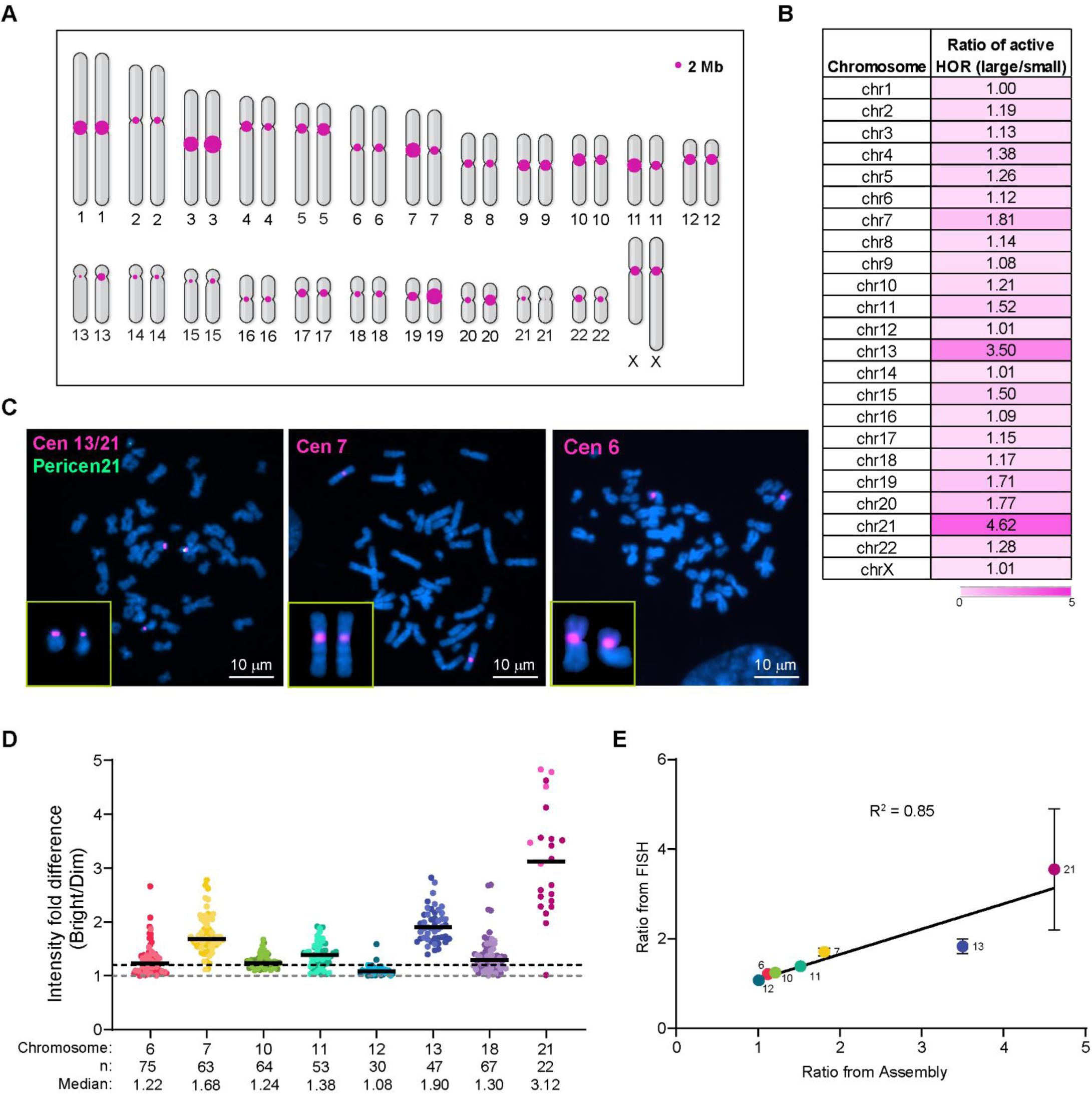
Quantitative FISH detects homolog-specific centromere array size variation. **A.** RPE-1 karyotype showing active α-satellite centromeric arrays for all chromosomes. Circle size is scaled with centromeric array size as determined by the T2T genome assembly of RPE-1. **B.** Ratios of the sizes of the active centromeric higher order repeat (HOR) between homologous chromosomes determined from the genome assembly are shown. Higher value (darker color) indicates greater difference in centromeric array sizes between homologous chromosomes. **C.** FISH on chromosome spreads detects active α-satellite centromeric arrays on chromosomes 6, 7, 13 and 21 in RPE-1 cells. Chromosomes 13 and 21, which have similar array sequences, are distinguished by a pericentromeric probe towards chromosome 21. Insets show a pair of homologous chromosomes with their centromere FISH signals. **D.** Fold difference in centromere FISH signal intensity is shown from homologous chromosome pairs in RPE-1 cells. Bar denotes the median. Fold-change of 1 denotes no difference in centromeric array size between homologs. Chromosomes with median higher than 1.2-fold (dotted line) indicate homologs with detectable size differences. **E.** Correlation is shown for fold-difference in homologous centromere sizes as determined by qFISH assay versus that obtained from the RPE-1 T2T genome assembly.

To replicate the observations made through inspecting the RPE-1 genome assembly, we pursued qFISH in chromosome spreads isolated from RPE-1 cells. We assessed a subset of chromosomes for which unique FISH probes were commercially available that target only active alpha-satellite repeat arrays (Figure 1C, Figure S1A, Table S1). To distinguish chromosome 13 from 21, which share nearly identical centromeric sequence, we used an additional probe targeting pericentromeric repeats present only on chromosome 21 (Figure 1C). Further, to reliably identify homologous pairs of chromosome 10, we employed a chr10 whole chromosome paint in addition to centromere FISH probes (Figure S1A). Thus, we queried 8 homologous chromosome pairs in RPE-1 cells (chr 6, 7, 10, 11, 12, 13, 18, and 21), and detected 4 (chr 7, 10, 11, and 13) with greater than 1.3-fold variation in centromere array size (Figure 1D). Variably sized homologous centromeres appeared as a pair of bright and dim FISH signal intensities. While an extreme size variation was visually observed between chr 21 homologs, the fold variation could not be reliably quantified through FISH on metaphase chromosomes because the smaller centromere 21 array only spans 321 kilobases in RPE-1 and is challenging to detect (Figure 1D). Using this methodology, we also queried 6 homolog pairs (chr 6, 7, 10, 11, 12, 18) in diploid HCT 116 cells, a cell line derived from colorectal carcinoma and observed greater than 1.3-fold difference in centromeric array sizes on two pairs (chr 6 and 12) (Figure S1B, D). To further validate our methodology, we performed qFISH on chromosome spreads from homozygous diploid HAP1 cells (Figure S1C). As expected, for all the chromosome pairs assessed (chr 6, 7, 10, 11, 12 and 18), intensity ratios ranged from 1.1 – 1.2, indicating minimal variation between homologous pairs (Figure S1E). The subtle variations in fluorescence in HAP1 cells could arise from minor genomic instability or depict experimental noise for the assay. We conclude that variation in centromeric array size beyond 1.2-fold can be reliably detected by our qFISH assay. Thus, we were able to empirically define three categories of centromeric array size on homologous chromosomes: 1) high size differences (chr 7, 13, 21 in RPE-1 and chr 6 in HCT 116), 2) moderate size differences (chr 10, 11, and 18 in RPE-1 and chr 12 in HCT 116) or 3) differences below our reliable detection limit (chr 6 in RPE-1 and chr 7, 10, 11 and 18 in HCT 116 cells). Hereafter we will refer to the third case as “undetectable.”

To confirm that the differences in FISH intensity ratios reflect the variation in length (kb) of homologous centromere pairs in RPE-1, we plotted centromeric array length ratios from qFISH versus the RPE-1 genome assembly ^26,27^ (Figure 1E). We found a close correlation (R^2^=0.85) (Figure 1E)^27^, except for centromere 18 that may have been assembled inaccurately, and we therefore excluded it from further analysis. This indicates that our qFISH assays accurately reflect differences in centromere size between a pair of homologous chromosomes.

We next queried if centromeric size differences between homologs can be reliably detected through different cell cycle phases. For this, we synchronized RPE-1 cells at G1 and G2 phases of the cell cycle (Figure S2A). We performed qFISH to detect a pair of signals for centromeres 6, 7, 10, 11, 13 and 22. The fold difference in intensities of the bright and dim array for each centromere remained invariant during the various phases of the cell cycle (Figure S2B), further confirming that qFISH is a robust method to detect size differences between homologous centromeric arrays.

Taken together, these results show that the differences in signal intensities identified by centromere qFISH reflect genetic differences in array size across homologous chromosomes. Thus, we have established a robust quantitative qFISH assay to distinguish the relative size of homologous centromeres in diploid heterozygous cells.

### Chromosome segregation fidelity at homologous centromeres with array size variants

We next asked how array size impacts centromere function during mitosis. As one readout of function, we investigated how the segregation of sister chromatids depends on centromeric array size by comparing homologous chromosomes. We synchronized RPE-1 cells in G2 phase and allowed one round of chromosome segregation followed by G1 arrest (Figure S2C). We then performed centromere qFISH for several chromosomes and proceeded with high-throughput imaging. When sisters segregate accurately, we expect two centromeric foci in each G1 nucleus (Figure S2C). Missegregation events, on the other hand, manifest as one or three foci. While we expected to find at least a few incorrect chromosome segregation events using our high-throughput imaging approach, baseline chromosome missegregation rates in RPE-1 cells were low and insufficient to score for centromere-specific biases (Figure S2D). Aneuploid nuclei with three centromeric foci ranged between 1.5-3% (Figure S2D). However, drug-induced spindle checkpoint inhibition significantly increased chromosome missegregation events in RPE-1 cells (Figure S2D) ^21^. Aneuploid nuclei with three centromere foci ranged between 6.6-9.8% (Figure S2D). Hence, we examined chromosome missegregation events in RPE-1 cells by synchronization in G2 followed by spindle-checkpoint inhibition and arrest in telophase, where every pair of daughter telophase nuclei represent a single chromosome segregation event (Figure 2A).

**Figure 2.**
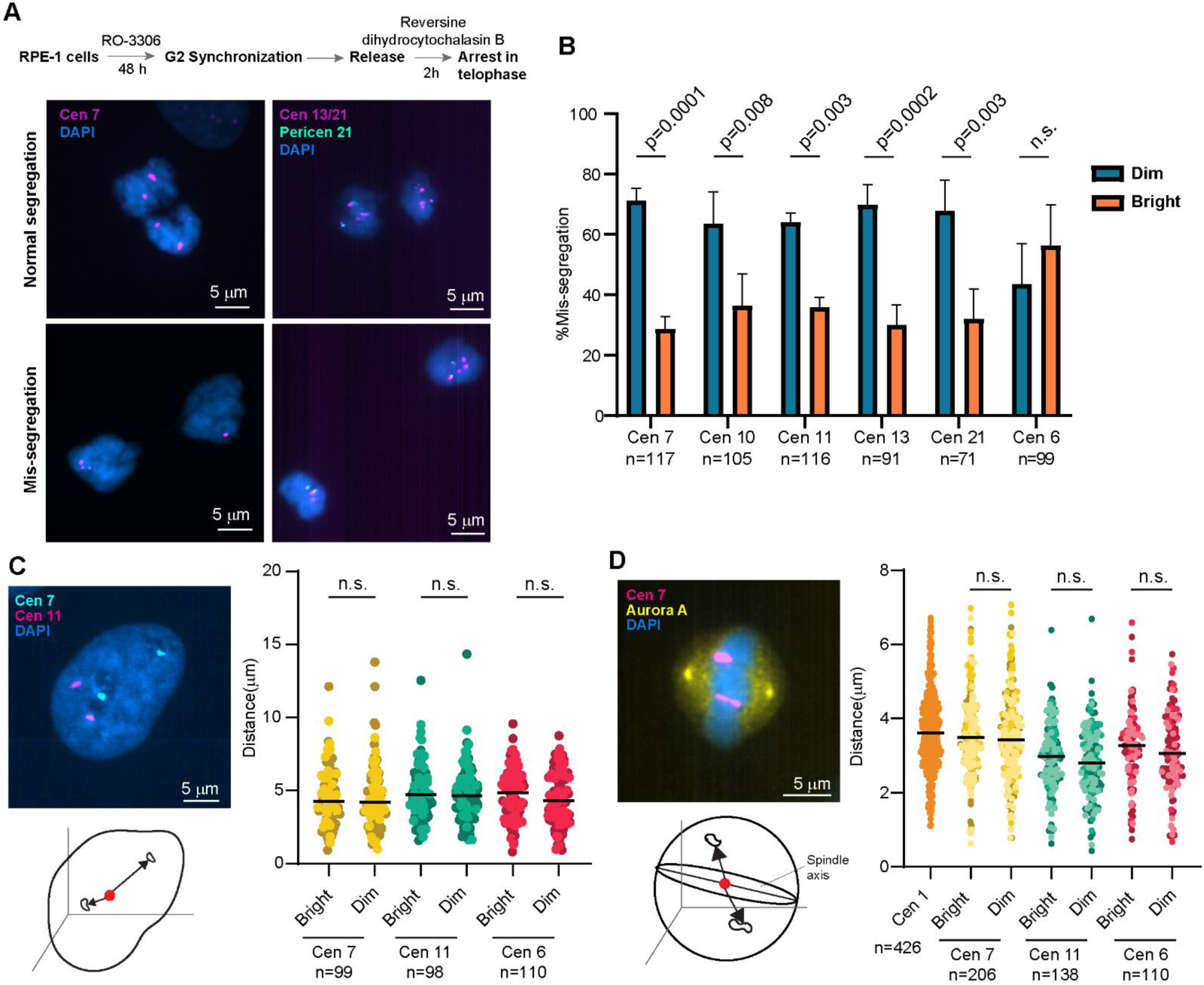
Chromosome missegregation events are enriched for the smaller centromere array. **A.** RPE-1 cells were synchronized in G2/M and released in medium containing reversine, a spindle checkpoint inhibitor and dihydrocytochalasin B, a cytokinesis inhibitor. Every pair of telophase nuclei represent one chromosome segregation event. Top panel shows a normal chromosome segregation event with one bright and one dim centromere signal in each daughter from sister chromatids that segregated correctly. Lower panel shows a chromosome missegregation event with three sisters that segregated to the left daughter nucleus and one that segregated to the right daughter. **B.** For all chromosomes assessed the homolog with the smaller centromere array (dim) has a higher rate of missegregation. Chromosome 6 homologs with no detectable difference in centromere array size do not show this bias. n denotes the number of chromosome missegregation events from three independent biological replicates. Error bar denotes S.D. Statistical significance tested by Chi-square test. **C.** 3D distances of bright and dim centromeres from the center of the nucleus in G2 synchronized cells were assessed. Bars denote median, n denotes number of centromeres assessed from three biological replicates. Significance tested by Wilcoxon paired signed rank test. **D.** Cells were synchronized in G2/M and then released and arrested in metaphase. ImmunoFISH was performed with centromere FISH probes and Aurora A staining to mark the centrosomes. 3D distance of centromere homologs from the plane connecting the spindle poles was determined. Bars denote median distance, n denotes number of cells assessed from three biological replicates. Significance was tested by Wilcoxon paired signed rank test.

We scored chromosome missegregation events where among a pair of daughter nuclei, one exhibited three centromeric foci and the other had only a single centromeric focus (Figure 2A). We queried several homologous chromosome pairs with distinct differences in centromere array sizes (chr 7, 10, 11, 13, 21) and one chromosome with undetectable difference (chr 6). Intensity measurements of centromeric foci revealed a pattern of asymmetry (Figure 2B). For chromosome 7, a pair exhibiting high difference (1.68-fold) in centromeric array size, the homolog harboring the dim centromeric array has a significantly higher chance of missegregation (71%) compared to the homolog with the bright centromeric array (29%) (Figure 2B). Similarly, we found that for chromosome 11, a pair exhibiting moderate difference (1.38-fold) in centromeric array size, the homolog harboring the dim centromeric signal has a significantly higher chance of missegregation (64%) compared to the homolog with the bright centromeric signal (36%) (Figure 2B). This pattern of biased missegregation of chromosomes harboring the dim centromere was also observed for several other homologous chromosome pairs exhibiting moderate to high variation in centromere sizes (chr 10, 13, and 21) (Figure 2B). However, for a chromosome pair with undetectable variation in centromere array size (chr 6), the array-size dependent bias in chromosome segregation pattern was not observed (Figure 2B). In this case centromeres were categorized as bright and dim based on slight differences in intensity. Thus, the longer active centromere haplotype segregates more accurately when the spindle checkpoint is compromised (Table S2).

Previously, the spatial localization of chromosomes in interphase and with respect to the spindle axis was shown to impact chromosome missegregation rates^24^. We asked if the observed bias in chromosome segregation could be explained by differences in spindle and chromosome positioning (Figure 2C, D). We assessed the positions of homologs of chromosomes 7 and 11. However, we did not observe any significant difference in distances of the bright and dim foci from the centroid of the nucleus for either chromosome pair (Figure 2C). Furthermore, bright and dim centromeres for both chromosome pairs are also equidistant from the spindle poles defined by Aurora A staining during metaphase (Figure 2D). This suggests that differences in chromosome position are not the mechanism for centromere size-based bias in chromosome missegregation.

Altogether, our study reveals that chromosome segregation fidelity is strikingly impacted by the size of the active centromere array. Thus, for pairs of homologous chromosomes with similar centromere HOR structures ^15,27^, the smaller centromeric array is more prone to chromosome segregation failure compared to the larger centromere array.

### Core centromere and kinetochore protein enrichment at centromeres with array size variation

Centromeric array size-based bias in chromosome segregation fidelity could originate from differences in core centromere and kinetochore protein enrichment. To this end, we screened the enrichment of the core centromere histone H3 variant, CENP-A, the centromeric DNA binding protein CENP-B, and several representative inner centromere proteins (INCENP, Aurora B), outer kinetochore subunits (KNL1, NDC80) and a fibrous corona subunit (CENP-E) on a pair of homologous centromeres with a size difference (Figure 3A). Although chromosomes 13 and 21 exhibit the largest differences in centromeric array size between homologs, acrocentric chromosomes have potential confounding factors like shape and rDNA content proximal to centromeres. Thus, we chose chromosome 7 for analysis, a metacentric chromosome with the third largest difference in centromeric array size in RPE-1 cells.

**Figure 3.**
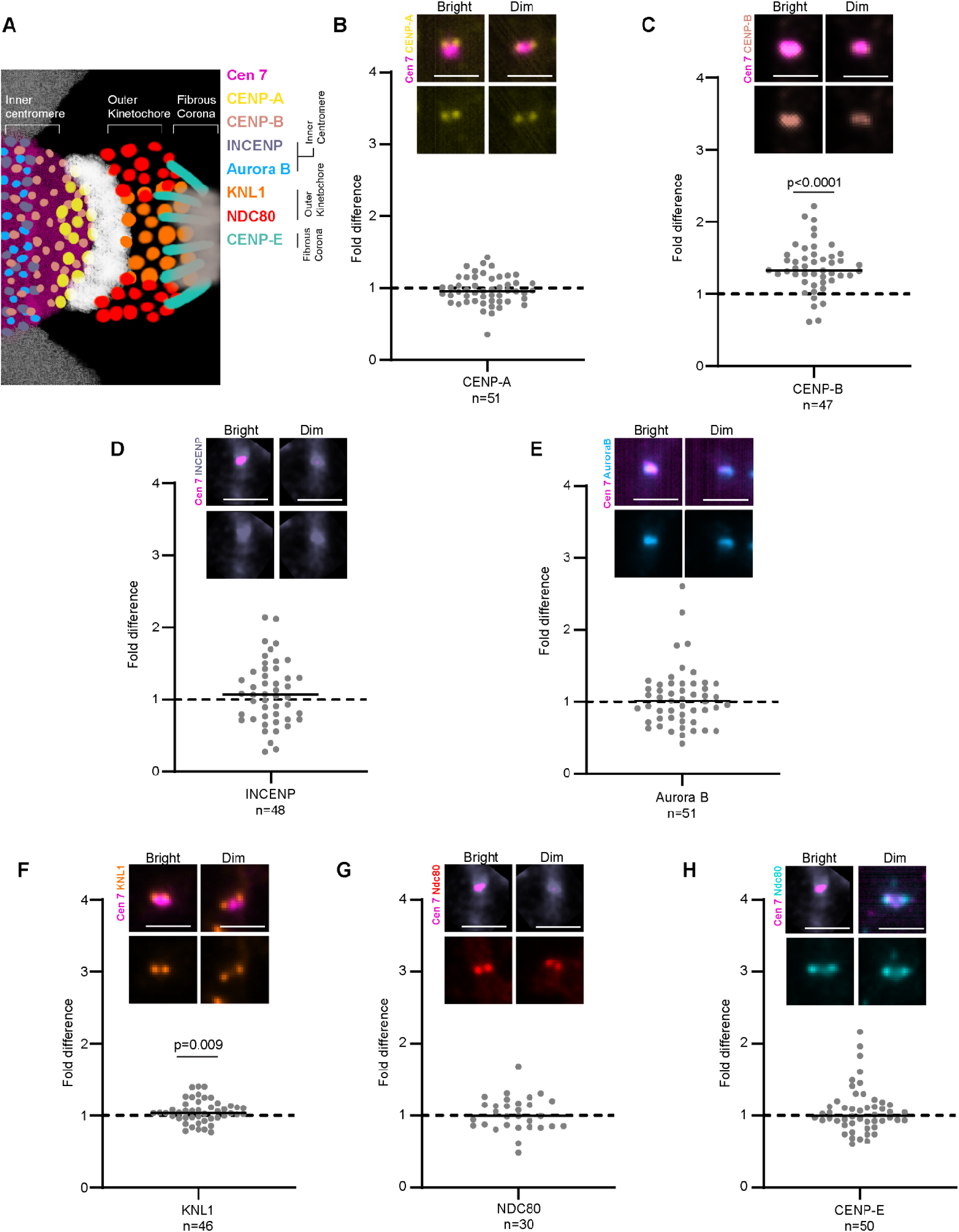
Homolog-specific enrichment of kinetochore proteins on centromere arrays. **A.** Cartoon of protein organization of the kinetochore. INCENP and Aurora B are inner kinetochore proteins, KNL-1 and Ndc80 are outer kinetochore proteins, and CENP-E is a fibrous corona protein. CENP-A is the centromere-specific histone H3 variant and CENP-B is a centromeric DNA binding protein. **B-H.** CENP-A, CENP-B, INCENP, Aurora B, KNL-1, Ndc80 and CENP-E levels on dim and bright centromeric arrays of chromosome 7 were determined by immunoFISH. Each dot represents the ratio of integrated density of centromeric and kinetochore protein enrichment on the bright over dim centromere 7 array from a single spread. Median fold change of 1 indicates no difference in respective protein enrichment. Insets show protein enrichment on the dim and bright centromere array from a pair of chromosome 7 homologs. Scale bars denote 1 μm. Statistical significance was determined using one sample t-test with a theoretical median of 1. n indicates the number of chromosome spreads assessed from three biological replicates for each protein.

To assess if levels of CENP-A underlie the array-specific difference in chromosome missegregation rates, we quantified CENP-A on centromere 7 arrays in RPE-1 cells from chromosome spreads by immunoFISH. Despite a 1.38-fold difference in array size, CENP-A levels showed no significant difference between the homologs, suggesting similar levels of the core centromeric chromatin that serves as the basis for kinetochore assembly (Figure 3B). We next quantified levels of CENP-B, a protein that binds to a 17 bp sequence present in alpha-satellite DNA. Levels of CENP-B have been shown to scale with centromere size, which we recapitulated (Figure 3C)^23^.

We further enquired if differences in kinetochore and inner centromere subunits could explain centromeric array size-based differences in chromosome missegregation rates. We found no significant difference in the levels of any examined kinetochore protein except KNL1, an outer kinetochore protein that showed 1.08-fold higher enrichment (p<0.01) on the larger centromere between the two homologs of chromosome 7 (Figure 3D-H). While statistically significant, the biological significance of a difference this small is unclear. Overall, core centromere and kinetochore proteins did not show a concerted and robust difference between homologous centromeres that would provide a satisfying explanation for functional differences. However, we have not exhaustively tested all proteins, all centromeres, and all conditions, and therefore acknowledge that centromere-specific levels of kinetochore proteins may influence centromere performance.

### Centromeric arrays exhibit distinct cohesion profiles during mitosis

Centromeres and pericentromeres maintain cohesion between sister chromatids until all chromosomes are bioriented on the metaphase plate and cells can proceed to anaphase. Failure to maintain cohesion results in precocious sister separation that can result in downstream chromosome segregation failure and aneuploidy. We next investigated cohesion at homologous centromeres as a potential size-based attribute. When cells are arrested in metaphase for prolonged periods with intact microtubules, cohesion can fatigue, leading to the separation of sister chromatids^30^. Cohesion fatigue is dependent on the cohesin complex but does not require its cleavage by separase^31^. To assess cohesion fatigue at centromeres, we first synchronized RPE-1 and HCT 116 cells in G2 phase and then arrested the cells in metaphase using an anaphase-promoting complex/cyclosome (APC/C) inhibitor (Figure 4A, Figure S3A). We observed two main centromeric conformations in metaphase-arrested cells (Figure 4B, Figure S3B). Some centromeres exhibited a continuous, elongated signal that we interpret as cohered sister chromatids experiencing microtubule-dependent tension. The second conformation was two discrete spots that we infer are sister centromeric arrays that have lost cohesion. To quantify centromeric cohesion, we measured the distance between sister centromeres (inter-centromere distance) measured the length connecting the centroids of the intensity profile of the centromere FISH signal (Figure 4C). Increased inter-centromere distance indicates reduced cohesion^32^.

**Figure 4.**
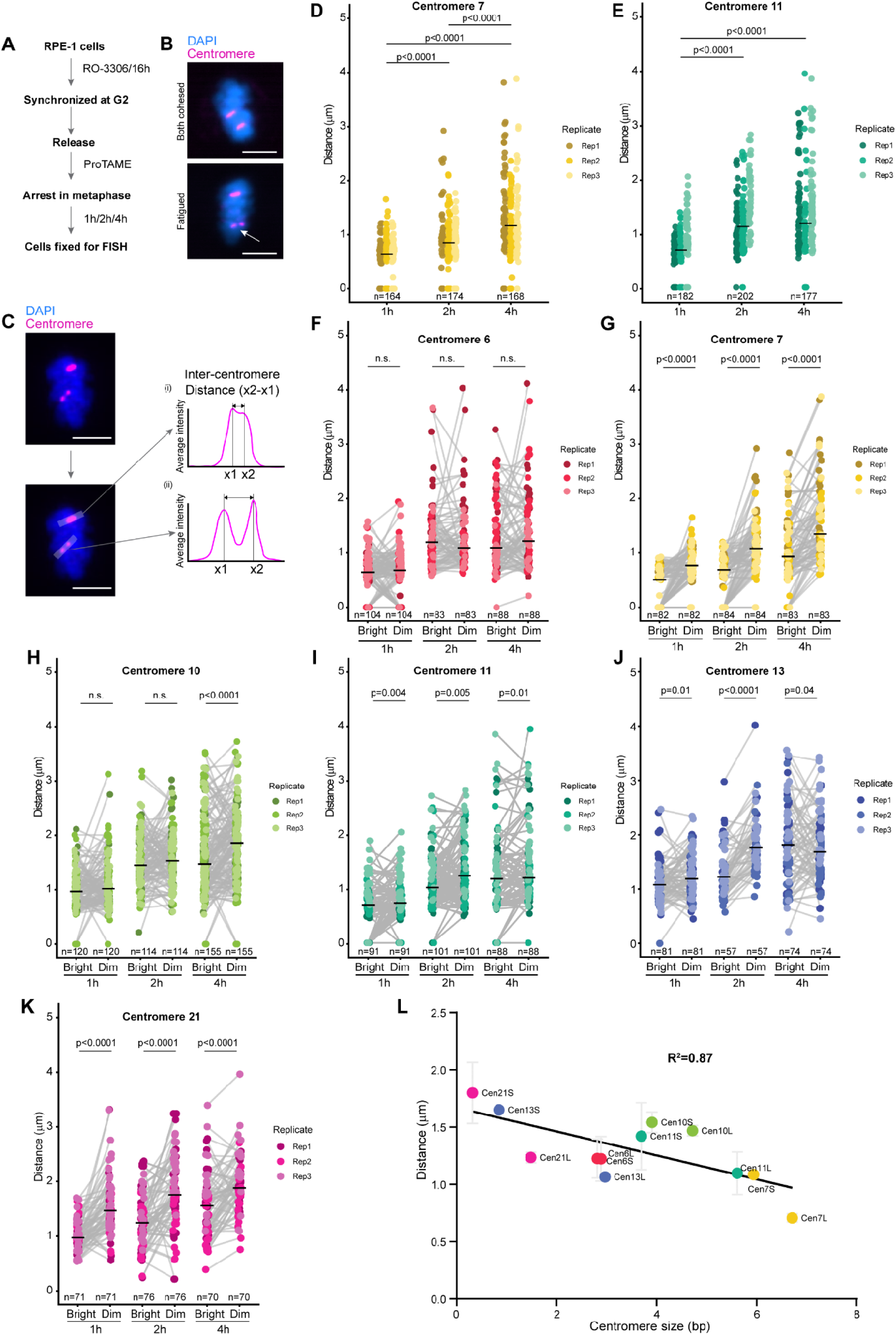
Centromere array size-based in cohesion in RPE1 cells upon prolonged metaphase arrest. **A.** RPE-1 cells were synchronized in G2/M and released in medium containing APC/Cdc20 inhibitor ProTAME for 1h, 2h, or 4h, then fixed for centromere FISH. **B.** Two centromere conformations were observed – cohesed (top panel both homologs show cohesed conformation) and fatigued (bottom panel, one centromere out of a homologous pair cohesed, other is fatigued). Scale bar denotes 5 μm. **C.** Strategy for assessing inter-centromere distances in RPE-1 cells arrested in metaphase after release from synchrony in G2 phase. Scale bar denotes 5 μm. **D-E.** Increasing inter-centromere distances for centromere 7 and 11 at 1h, 2h or 4h of metaphase arrest showing loss of cohesion as a function of time. Bars indicate median distance. Statistical significance tested by Kruskal-Wallis test. **F-K.** Pairwise comparison of inter-centromere distances at bright and dim centromeres at 1h, 2h and 4h of metaphase arrest from chromosomes 6, 7, 10, 11, 13, and 21. Dim centromeres show higher inter-centromere distances. Bars indicate median distance. Statistical test is Wilcoxon test. n denotes the number of cells assessed in every condition from three biological replicates. n.s. denotes not significant. **L.** Inter-centromere distances from 2h of metaphase arrest are plotted as a function of centromeric array size determined from the RPE-1 T2T assembly. S and L denote chromosome homologs with smaller and larger centromere versions, respectively. Error bars denote S.D.

As expected, with increasing time in metaphase arrest, the inter-centromere distance steadily increases for all centromeres assessed in RPE-1 cells (Figure 4D, E, Figure S3C-E) and HCT 116 cells (Figure S3F-H), consistent with previous work demonstrating that cohesion is lost upon prolonged mitotic arrest^30,31^. The median separation distance for centromeres after two hours of mitotic arrest is higher in HCT 116 cells than that of comparable centromeres in RPE-1 cells, indicating that centromeres in HCT 116 cells are more susceptible to cohesion fatigue upon prolonged mitotic arrest. A possible explanation for this could be lower mitotic chromatin compaction in HCT 116 compared to RPE-1 cells^33^.

We next compared cohesion fatigue between pairs of homologous centromeres (chr 6, 7, 10, 11, 13, and 21 in RPE-1 and chr 6, 12, 18 in HCT 116 cells) (Figure 4F-K, Figure S3I-K). We found that all homologous chromosome pairs exhibiting greater than 1.3-fold difference in centromere array size (chr 7,11,13, 21 in RPE-1 and chr 6 and 12 in HCT 116) show significantly higher inter-centromere distance for the dim compared to the bright centromere, at every assessed timepoint (Figure 4G, 4I-K, S3I,J). The smaller centromere array for chr 10 in RPE-1 cells showed significantly higher inter-centromere upon 4 hours of mitotic arrest, showing that this relationship is true even for homologous pairs with moderate size difference (1.2-fold) during longer arrest times (Figure 4H). However, for chromosome homologs with an undetectable difference in size (chr 6 in RPE-1 and chr 18 in HCT 116), both centromeres exhibited similar inter-centromere distance at all assessed timepoints (Figure 4F, S3K). These results suggest that between a pair of homologous chromosomes, short centromeric arrays are more prone to cohesion fatigue compared to long arrays, and the effect is more pronounced with larger size differences.

Our ability to identify six individual centromere pairs in RPE-1 cells based on their signal intensities allowed us to build a profile of centromere cohesion fatigue at 12 individual chromosomes relative to each other (Figure 4L). We examine the correlation between centromere array length as determined by the RPE-1 assembly, and cohesion fatigue cross several chromosomes in RPE-1 cells identified by their large (chr 6L, 7L, 10L, 11L, 13L and 21L) or small (chr 6S, 7S, 10S, 11S, 13S and 21S) centromeres. We found that array size negatively correlated with cohesion fatigue. However, the correlation is imperfect and additional factors likely contribute to centromeric cohesion (Figure 4L).

Taken together, these results show that between a pair of homologous centromeres, the centromere with the smaller array size is more prone to cohesion fatigue than the larger centromere. Moreover, the cohesion fatigue profile roughly scales with the difference in array size between homologous pairs of centromeres, suggesting cohesion is a size-dependent feature of centromeric arrays. When differences in centromere size are not robustly detected by FISH, we cannot detect appreciable differences in cohesion fatigue between homologs. Finally, size-based differences in cohesion fatigue appear agnostic of alpha-satellite sequence composition, as we observe these effects at centromeres with diverse HOR structures^34^. Future work will be needed to understand underlying chromosome-specific differences in cohesion.

### Mechanisms underlying size-based bias in centromere array cohesion

Given the scaling of cohesion fatigue with centromeric array size, we were motivated to understand the mechanisms that underlie more robust cohesion at larger centromere arrays. Multiple factors mediate cohesion at centromeres including the cohesin complex and DNA catenations regulated by topoisomerase 2A^35,36^. Further, CENP-B has been previously shown to impact centromeric cohesion and compaction^37,38^. To gain an understanding of factors that could generate the size-based bias in centromere cohesion, we targeted the cohesin complex, topoisomerase 2A and CENP-B.

### Role of CENP-B

In our current study and previous studies, CENP-B protein is found to scale with centromere array size (Figure 3). Thus, CENP-B is an attractive candidate for mediating size-based bias in centromere cohesion. We depleted *CENPB* in RPE-1 cells for 48 hours using siRNAs and synchronized the cells in G2 (Figure 5A). We then released cells from G2 arrest into metaphase arrest for 2 hours, keeping microtubules intact. Next, we assessed inter-centromere distance from two sets of homologous centromeres (cen7 and 11) (Figure 5B). Upon CENP-B depletion, we found a significant increase in inter-centromere distance for both chromosomes indicating an overall loss of cohesion (Figure 5C, D). We then asked if the size-based bias in centromere cohesion is lost or maintained upon CENP-B depletion. Control siRNA treated cells recapitulated our previous result and the bright centromeres of both chromosomes exhibited lower inter-centromere distance and better cohesion (Figure 5E, F). The bright centromere of chromosome 7 maintained higher cohesion even after CENP-B depletion (Figure 5E). However, upon CENP-B depletion, cohesion at both bright and dim centromeres was no longer significantly different for chromosome 11 (Figure 5F). This indicates that a homologous chromosome pair with high variation in centromeric array size maintains the bias in centromere cohesion upon CENP-B loss, whereas the cen11 chromosome pair with moderate variation is dependent on CENP-B for cohesion bias, which could be a size or sequence effect.

**Figure 5.**
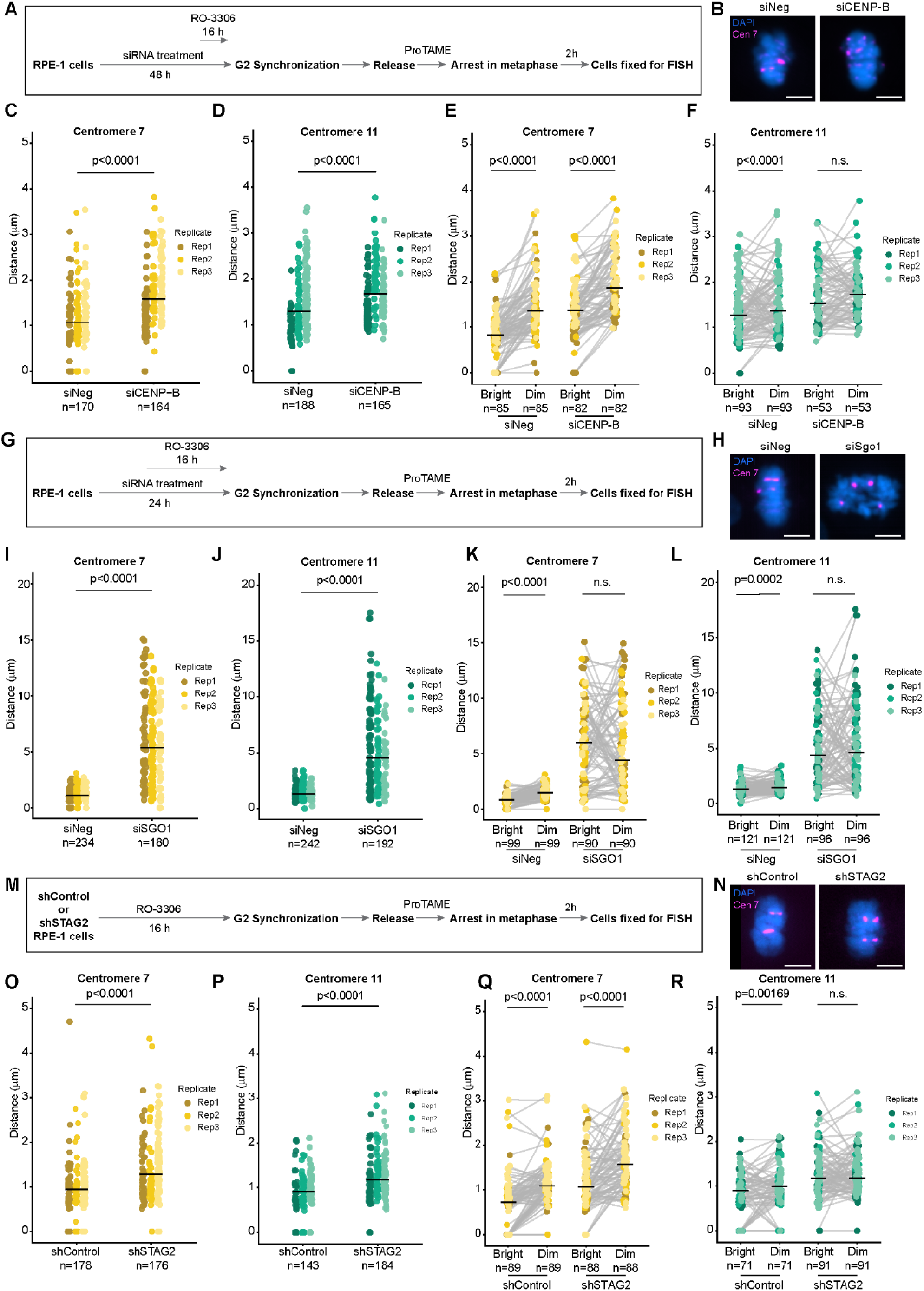
Cohesin complex determines centromere array-size based differences in cohesion. **A.** RPE-1 cells were depleted of CENP-B by siRNA treatment and synchronized in G2/M, after which they were released into medium containing ProTAME. **B.** Representative images of centromere FISH performed on control siRNA (siNeg) and *CENP-B* siRNA (siCENP-B) treated cells arrested in metaphase for 2 hours. Control cell shows one cohesed and one fatigued centromere. siCENP-B cell shows both homologs fatigued. **C-D.** siCENP-B cells show significantly higher inter-centromere distances for both chromosome 7 and 11. Statistical significance tested with Kruskal-Wallis test. **E-F.** Pairwise comparison of inter-centromere distances at bright and dim centromeres of chromosome 7 and 11 in siControl and siCENP-B cells. Dim centromeres out of a homologous pair show significantly higher distance in both siControl and siCENP-B treated cells. Statistical significance tested by Wilcoxon test. **G.** RPE-1 cells were depleted of *SGO1* by siRNA treatment and synchronized in G2/M, after which they were released into medium containing ProTAME. **H.** Representative images of centromere FISH performed on control siRNA (siNeg) and *SGO1* siRNA (siSGO1) treated cells arrested in metaphase for 2 hours. Control cell shows one cohesed and one fatigued centromere. siSGO1 cells show massive loss of sister chromatid cohesion and both centromere 7 homologs fatigued. **I-J.** Inter-centromere distances are significantly higher in siSGO1 treated cells compared to controls for both chromosomes 7 and 11. Statistical significance tested by Kruskal-Wallis test. **K-L.** Pairwise comparison of inter-centromere distances at bright and dim centromeres of chromosome 7 and 11 in siControl and siSGO1 cells. Dim centromeres out of a homologous pair show significantly higher distance in control cells. Inter-centromere distances between dim and bright centromeres are no longer significant upon SGO1 depletion. Statistical significance tested by Wilcoxon test. **M.** RPE-1 cells with constitutive STAG2 knockdown (shSTAG2) or empty vector (shControl) were synchronized in G2/M and then arrested in proTAME for 2 or 4 hours. **N.** Representative images show one cohesed and one fatigued centromere in shControl cells and both centromeres fatigued in shSTAG2 cells. **O-P.** shSTAG2 cells show significantly higher inter-centromere distance compared to shControl cells for chromosomes 7 (4 h) and 11 (2h). Statistical significance tested by Wilcoxon test. **Q-R.** Pairwise comparison of inter-centromere distances at bright and dim centromeres of chromosome 7 and 11 in shControl and shSTAG2 cells. shControl cells recapitulate the previous results that the bright arrays of centromere 7 and 11 show better cohesion than their dim counterparts. Upon STAG2 depletion, centromere 7 pairs maintain the size-based differences in cohesion, while centromere 11 pairs show no difference in cohesion. Statistical significance tested by Wilcoxon test. Bars denote median inter-centromere distance. n denotes the number of cells assessed for every condition from 3 biological replicates. n.s. - not significant. Scale bars represent 5 μm.

### Role of cohesin complex

Next, we determined the contribution of the cohesin complex in mediating size-based bias in centromere cohesion. Cohesin is a multi-subunit complex and its activity at centromeres can be modulated by targeting individual subunits of the complex or cohesin regulators. First, we depleted shugoshin-1 in RPE-1 cells (Figure 5G). Shugoshin-1 (SGO1) protects centromeric cohesin complexes from degradation during mitosis^39^. While we did not observe a significant difference in SGO1 enrichment on bright and dim centromere arrays of chromosome 7 (Figure S4A, B), we asked if complete deprotection of cohesin complexes by SGO1 depletion at centromeres relieves the array size-based bias in cohesion. As expected, SGO1 depletion caused a significant loss in sister chromatid cohesion at all centromeres assessed (Figure 5H-J). Furthermore, SGO1 depletion equally impacted both bright and dim centromere pairs of chromosome 7 and 11(Figure 5K, L). This shows that the cohesin complex is responsible for the size-based bias in centromere cohesion, as upon its complete removal, both large and small centromeres are equally susceptible to cohesion fatigue.

Cohesin exists in two distinct versions in human somatic cells – one that contains STAG1 and one containing STAG2. Depletion of the STAG1 subunit decreases telomeric cohesion whereas depletion of the STAG2 subunit results in loss of centromeric cohesion^40^. We next asked how loss of STAG2-cohesin impacts sister chromatid cohesion at homologous centromeres with measurable size differences. We constitutively depleted *STAG2* in RPE-1 cells using stable shRNA integration and arrested cells in metaphase to allow cohesion fatigue (Figure 5M). Compared to control RPE-1 cells, STAG2 depleted cells (shSTAG2) show significantly higher inter-centromere distances at both chromosome 7 and 11 (Figure 5N). This recapitulates previous studies in other cell lines demonstrating that STAG2 preserves centromeric cohesion.

We next assessed cohesion as a function of centromeric array size for chromosomes 7 and 11. Consistent with our previous experiments, we observed significantly greater inter-centromere distances at the dim centromere compared to the bright centromere of chromosomes 7 and 11 in control cells (Figure 5O, P). For chromosome 7, the bias in cohesion fatigue between the dim and bright centromere arrays is maintained in the absence of cohesin-STAG2 (Figure 5Q). In contrast to chromosome 7, cohesion at bright and dim centromeres is no longer significantly different for chromosome 11 upon cohesin-STAG2 depletion (Figure 5R). Therefore, the size-based centromeric cohesion profile displays varying dependency on cohesin-STAG2. Like CENP-B, this differential dependence on STAG2 could be based on size (high versus moderate), sequence (HOR structure), or a combination thereof.

### Role of catenations

In addition to the cohesin complex, DNA catenations at centromeres play an important role in preserving cohesion^35^. We inspected catenations as a size-dependent feature of centromeres that may scale with DNA length. After chromosome biorientation at the mitotic spindle, catenations are removed by the enzyme topoisomerase 2A (TOPO2A), ensuring accurate chromosome segregation. We hypothesized that retaining catenations by inhibiting TOPO2A may eliminate the bias in centromeric cohesion between large and small centromeres. Although the total enrichment of TOPO2A at bright or dim centromeres of chromosome 7 was not different (Figure S4C, D), centromeres could be differentially impacted by TOPO2A inhibition. Many TOPO2A inhibitors are commercially available but most arrest cells in prometaphase^41^. Hence, we synchronized RPE-1 cells in G2 and arrested them in metaphase with intact microtubules in the presence or absence of the TOPO2A inhibitor merbarone^42^ (Figure 6A). TOPO2A inhibition and retention of catenations, as evidenced by PICH staining during metaphase (Figure 6B), significantly reduced inter-centromere distances for all chromosomes assessed (chr 7 and 11), minimizing cohesion fatigue (Figure 6C, D). We next asked if catenation preservation abolished array size-based bias in cohesion. Contrary to our hypothesis, we found that for chromosome 7 pairs, the dim centromere was still more susceptible to cohesion loss than the bright centromere, regardless of TOPO2A inhibition (Figure 6E, F). Thus, enhanced catenation at larger centromeres does not seem to be the driving mechanism for better cohesion, at least under the conditions tested.

**Figure 6.**
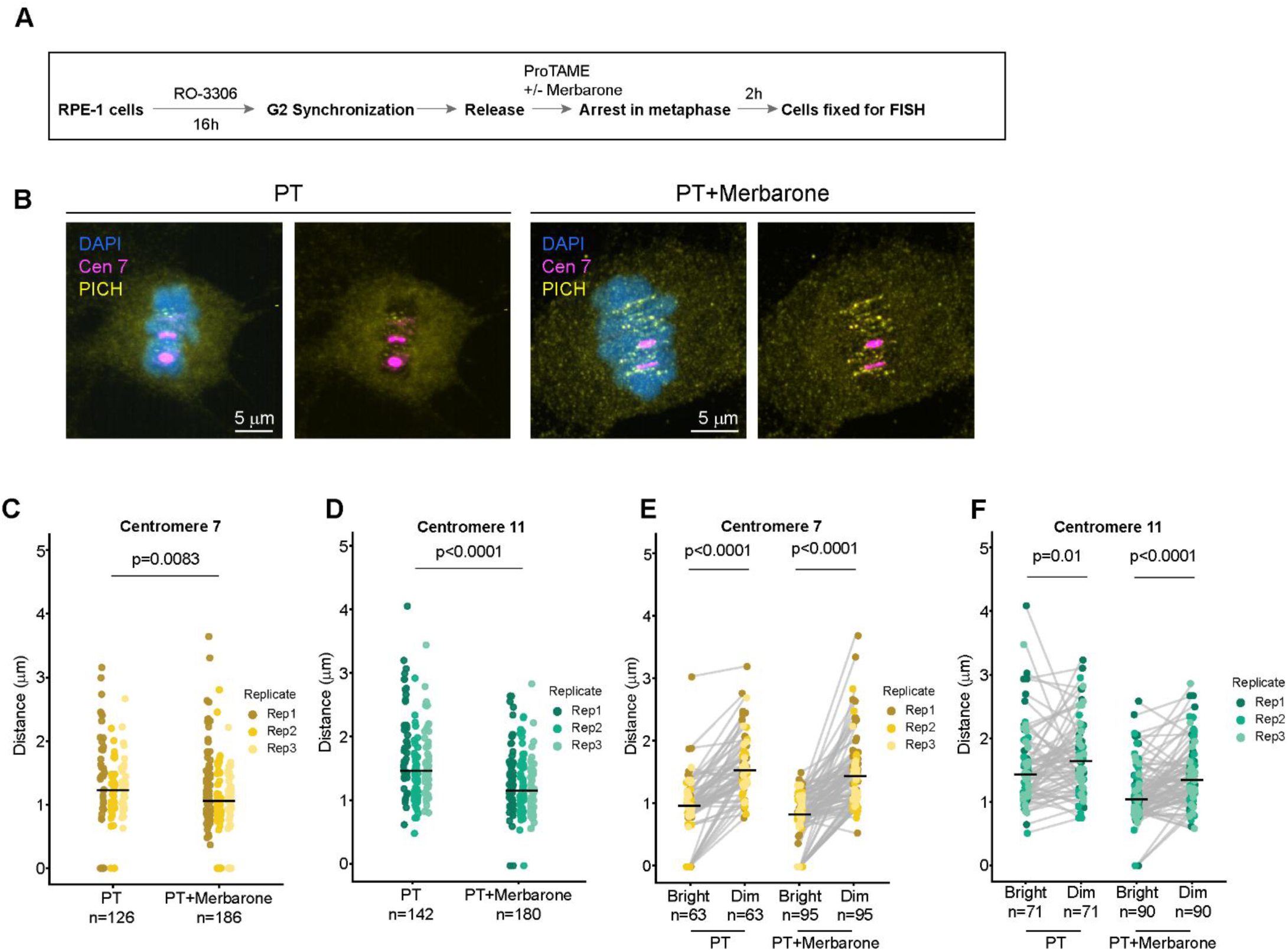
Topoisomerase 2A inhibition retains array size-based bias in centromeric cohesion. **A.** RPE-1 cells were synchronized at G2/M and released in medium containing topoisomerase 2A inhibitor merbarone and APC/cdc20 inhibitor ProTAME (PT). Cells were arrested in metaphase for 2 hours and fixed. **B.** Representative immunoFISH images show cells treated with only ProTAME, or ProTAME with merbarone. PICH staining is enhanced in cells with merbarone treatment indicating effective TOPO2A inhibition. **C-D.** Inter-centromere distances are significantly reduced upon topo 2A inhibition at both chromosomes 7 and 11. Statistical significance tested by Kruskal-Wallis test. **E-F.** Pairwise comparison of inter-centromere distances at bright and dim centromeres of chromosome 7 and 11 in control and merbarone treated cells. Dim centromeres out of a homologous pair show significantly higher distance in control cells. Inter-centromere distances remain significantly higher at dim centromeres of chromosome 7 and 11, upon TOPO2A inhibition. Statistical significance tested by Wilcoxon test.

Collectively, these results confirm that the cohesin complex is broadly required for centromeric cohesion and further suggests centromeric cohesin mediates the array size-based bias in cohesion at homologous centromeres of unequal sizes. Cohesin-STAG2 and CENP-B are partially responsible for the size-based bias at specific chromosomes. Catenations did not impact centromere array size-based bias in cohesion for the centromeres studied. Future investigations will reveal how SMC complexes, heterochromatin, and other centromere associated proteins are differentially enriched or positioned at larger centromeric arrays to form pericentromeres that are more resistant to spindle forces.

Our results strongly suggest that aspects of mitotic centromere function, including sister chromatid cohesion and segregation fidelity, can be influenced by centromeric array length, especially when centromere function is challenged. Importantly, the length effects are observed across centromeres with different alpha satellite composition and in different cell lines, suggesting the effect depends on centromere size rather than sequence differences.

## Discussion

Recent long-read sequencing and assembly efforts have made immense progress in identifying genomic variation at human centromeric arrays. For homologous chromosomes across individuals, the centromeric higher order repeat (HOR) sequences tend to be highly similar, but the size of the active HOR can vary up to 21.5-fold^43^. Herein we have capitalized on these advances to address the consequences of centromeric array size variation and propose a model to explain how these differences impact mitotic performance. Through our qFISH assays, we identified multiple pairs of homologous chromosomes in RPE-1 and HCT 116 cell lines that varied substantially in centromeric array size and documented that the smaller centromeric array is more prone to cohesion loss and predisposes the homolog to aneuploidy. Our results are consistent with previous studies on heterologous chromosomes, that suggested that between chromosomes of comparable sizes, larger centromeric arrays are better at preserving cohesion and have a lower propensity to missegregate^21–23^. However, these studies were limited by the simultaneous analysis of centromere size averaged across homologs, unresolved chromosome-specific sequence differences in α-satellite DNA composition, and the lack of distinction between the active and inactive α-satellite arrays.

By comparing multiple pairs of homologous chromosomes, our study disentangles size effects of centromere arrays from potentially confounding α-satellite sequence variation. To our knowledge, this study presents the largest to date functional assessment of chromosome cohorts stratified by centromere array size and provides compelling evidence that centromere size can influence sister chromatid cohesion and chromosome segregation fidelity. Gao et al, 2025 assessed hundreds of genomes and found centromere array sizes vary by 1.5-fold on average between homologous chromosomes^43^. As a further example, homologs of chromosomes 3 and 21 in the HG002 genome exhibit similar size variation as homologs of chromosome 7 in RPE-1 cells^44^. Therefore, we speculate that size-based functional differences will be pertinent in most genomes.

For all homologous pairs of chromosomes assessed in this study, we uncover a relatively higher propensity of missegregation at smaller centromere arrays when the spindle checkpoint is abrogated. This reinforces findings from previous studies in RPE-1 cells where higher overall centromere to chromosome size ratio results in better chromosome segregation fidelity^23,25^. Paradoxically, a study in immortalized Indian muntjac fibroblasts demonstrated that a chromosome with a 2-fold larger array is more prone to merotelic attachments and missegregates at a higher rate, although it is better at chromosome congression^22^. We speculate that this paradox can be explained by size-based differential CENP-A and kinetochore enrichment at larger muntjac centromeres promoting merotely, an attribute that is not observed at RPE-1 chromosome homologs described here. CENP-A distribution is more uniform across human chromosomes, although Y-chromosomes and neocentromeres exhibit lower levels^45,46^. In our study, we observed some variation in kinetochore protein enrichment across chromosomes within single spreads. In our calibrated studies, the amount of CENP-A per spread varies by an average of 1.7-fold (+/-0.15 S.D.), CENP-B varies an average of 2.8-fold (+/- 0.3 S.D.), and kinetochore subunits NDC80 and KNL1 vary an average of 1.8-fold(+/- 0.17 SD) and 1.9-fold(+/- 0.12 SD), respectively.

However, we did not uncover significant differences in enrichment of kinetochore proteins when comparing cen7 homologs. Moreover, the variation in protein level is lower than the difference in array size across the RPE1 genome. Thus, kinetochore heterogeneity may be a centromere attribute that can influence cohesion and segregation, but more investigation will be required to examine this possibility.

We propose a working model to explain functional differences in centromeres based on size. In our model we illustrate homologous chromosomes with relatively smaller and larger centromere array sizes (Figure 7). Sister chromatid cohesion during mitosis is likely achieved by a combination of factors including, but not limited to, the cohesin complex, which is enriched at pericentromeres^47^, DNA catenations that exist throughout chromosome arms^7,35,48^, and CENP-B dependent DNA structure^38^. Furthermore, some centromeres are bipartite, another structural feature that may have implications for size-based centromere strength^49^. Our work reconfirms the critical role of cohesin, DNA catenations, and CENP-B as master regulators of centromeric cohesion, affecting all centromeres. However, out of the three mechanisms of cohesion that we investigated, cohesin complex-mediated cohesion was the most significant determinant of array size-based differences in centromere cohesion, as uncovered by depletion of the centromeric cohesin protector shugoshin.

**Figure 7.**
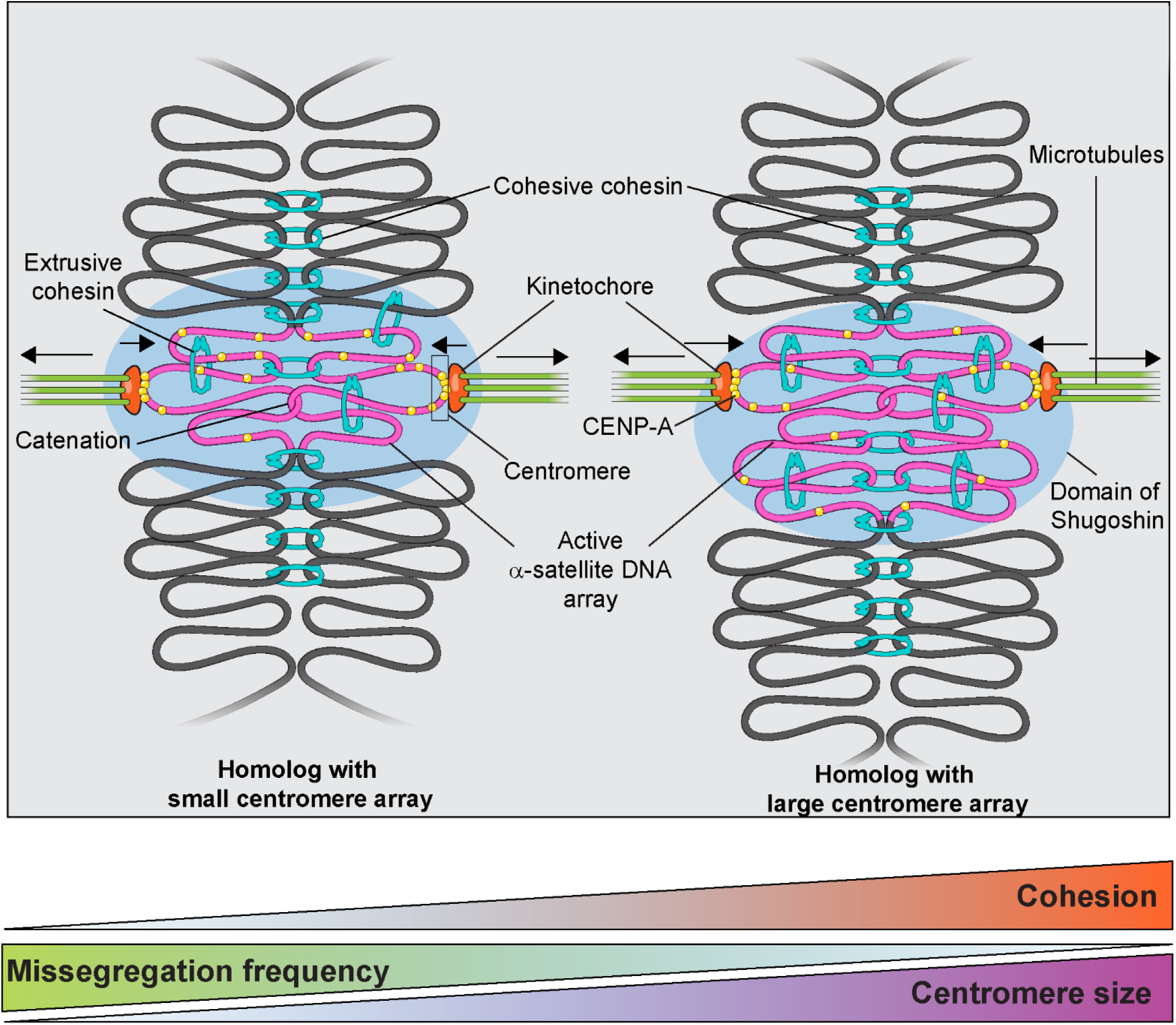
Model depicting the structural attributes of centromere arrays underlying size-based differences in performance. During early metaphase, sister chromatids remain cohesed largely by the cohesive cohesin complex (cyan) and catenations. Chromatin at the primary constriction is organized into loops by loop-extruding cohesin (cyan) and other SMC complexes (omitted for simplicity). Active α-satellite DNA array (pink) is epigenetically marked by CENP-A (yellow circles), but CENP-A is most enriched at the core centromere (open black box), that contacts the kinetochore complex (orange) and kinetochore microtubules (green). Any DNA outside of the active α-satellite is shown in gray. Cohesin is removed from chromosome arms by prophase pathway but retained at the primary constriction (spanning pink and gray) during metaphase. Kinetochore microtubules exert outward force while the core- and peri-centromere provide resistive inward force (black arrows). In our model, the smaller active α-satellite array (pink) occupies less space in 3D, forms fewer chromatin loops, and recruits fewer molecules of cohesin, compared to the larger array. The magnitude of resistive force (inward arrows) generated by the smaller α-satellite array directly adjacent to the core centromere is significantly lower compared to the larger array. Cohesin complexes are protected by a large domain of shugoshin enrichment (light blue cloud) at core and pericentromeric regions. Depletion of shugoshin results in loss of cohesin complexes, and both large and small arrays become equally susceptible to kinetochore microtubule forces. Cohesion scales positively with active α-satellite array size while missegregation scales negatively and is uncovered under perturbed cell cycle conditions.

Cohesion is achieved at pericentromeric domains through a combined action of cohesive and loop-extruding cohesin complexes^50^. Alleviation of the array size-based differences in centromere cohesion upon shugoshin-1 knockdown strongly implicates cohesin complexes in mediating this function. The similar levels of SGO1 on cen7 haplotypes points to differences in cohesin enrichment, as opposed to enhanced cohesin protection^51^, at larger centromeres. Based on our previous study, the smaller centromere array within a homologous pair occupies less three-dimensional volume relative to the larger array^47^. We speculate that the smaller centromere array harbors fewer cohesin molecules. Microtubule-dependent tension originates at the similarly sized kinetochore domain. As the tension extends to the flanking centromeric DNA, cohesin complexes progressively disengage, leading to cohesion fatigue. In our model, the number of cohesin complexes scales with centromeric array size, conferring array size-based differences in cohesion. Additional factors could operate independently or in conjunction with cohesin, where shorter centromeres confer lower centromere strength to counteract microtubule mediated tension. Factors contributing to strength may include condensin, heterochromatin, and checkpoint proteins.

Our study is significant because it demonstrates that centromere genotypes impact the fidelity of chromosome segregation. These findings have broad implications in the context of cancer, infertility, and other disorders characterized by aneuploidy. Prolonged delay in mitosis is common in cMyc overexpressing cancer cells, and cancer cells undergoing chemotherapy^52,53^. Furthermore, the spindle checkpoint is also often compromised in cancer cells and aneuploidy is a frequent occurrence. Centromeres must maintain sister chromatid cohesion during the prolonged meiotic arrest, and loss of sister chromatid cohesion increases with maternal age^54–56^. We speculate that an individual’s genetically inherited complement of centromeres could contribute to personalized patterns of chromosome segregation errors and aneuploidy. While our study provides strong evidence for a size-based mechanism regulating cohesion and segregation, the molecular mechanisms underlying these observations will require additional studies.

## Materials and Methods

### Cell culture

hTERT RPE-1 cells and derivatives were sourced from ATCC (CRL-4000) and cultured in DMEM:F12 medium (Gibco) supplemented with 10% FBS (Peak Serum Inc.) and 1X Glutamax (Gibco). HCT 116 cell line was a gift from Jan-Michael Peters (IMP, Vienna) and cultured in McCoy5A medium (Gibco) supplemented with 10% FBS. HAP1 cells were sourced from Horizon (C859) and cultured in Iscove’s medium supplemented with 10% FBS. HAP1 cells were sub-cultured up to 8 times enriching for diploid populations^57^. All cells were cultured at 37^0^C, 5% CO_2_ under humidity-controlled conditions. Cells were routinely tested for the absence of *Mycoplasma* contamination using PCR.

### Cell synchronization and pharmacological treatments

RPE-1 cells were synchronized in G1 using Palbociclib (1 μM, Selleckem, S1116). For loss of cohesion assays, cells were first synchronized in RO-3306 (10 μM, ENZO LIFE SCIENCES INC, ALX-270-463-M001) for 16 hours and released in medium containing ProTAME (20 μM, Tocris, 7734) for RPE-1 cells or MG-132 (10 μM, Sigma-Aldrich, 474791) for HCT 116 cells. For chromosome missegregation assays, RPE-1 cells were synchronized in G2-phase using RO-3306 for 16 hours, washed and released in complete medium containing the MPS1 inhibitor reversine (250 nM, Sigma-Aldrich, R3904) and actin inhibitor dihydrocytochalasin B (10 nM, Sigma-Aldrich, D1641). For topoisomerase 2A inhibition, cells were synchronized in G2-phase using RO-3306 for 16 hours, washed and released in complete medium containing ProTAME and merbarone (100 μM, Cayman Chemicals, 97534-21-9) for 2 hours. Cell cycle profiles were assessed by cytometry with propidium iodide staining (Figure S5).

### Gene knockdown

shControl and shSTAG2 cells were derived from hTERT RPE-1 cells by transducing the cells with Dharmacon GIPZ Lentiviral shRNA particles containing Empty vector shRNA control and *STAG2* shRNA (Cat. No. VGH5518-200288547, *STAG2* gene targeting sequence: 5’-TGGATCATGTCTTCATTGA-3’), respectively. Positive cells were bulk sorted using a BD FACSymphony S6 based on the presence of Turbo-GFP fluorescence. *SGO1* and *CENPB* were depleted in RPE-1 cells by ON-TARGETplus SMARTPool siRNA (Horizon, SGO1: L-015475-00-0010, *CENPB*: L-003250-00-0010) reverse transfection using RNAimax (Thermo Fisher, 13778075). Negative control siRNA was ON-TARGETplus Non-targeting Pool (Horizon, D-001810-10). All siRNAs were used at a final concentration of 40 nM. Knockdown efficiency was determined by Western Blotting of whole cell lysate (Figure S6). Primary antibodies were used as follows: rabbit anti-SA2 (Bethyl, A302-581A, dilution 1:1000), rabbit anti-SGO1 (Abcam, ab225948, dilution 1:500), rabbit anti-CENP-B (Abcam, ab25734, 1:1000). Loading controls used were: mouse anti-Actin (Abcam, ab6276, dilution 1:1000), mouse anti-GAPDH (Invitrogen, AM4300, dilution 1:10000), and rabbit anti-βTubulin (Cell Signaling, 2128, dilution 1:1000).

### Chromosome spread preparation

HAP-1, RPE-1 and HCT 116 cells were treated with KaryoMAX Colcemid solution (100 μg/mL) for 2-4 hours in complete medium to enrich mitotic cells. Cells were dissociated using TrypLE Express (Gibco) and collected by centrifugation at 1200 rpm. Cells were then treated with hypotonic 0.075 M potassium chloride (Gibco) hypotonic solution for 10 minutes at 37^0^C, pre-fixed with 3:1 methanol:acetic acid (v/v) and centrifuged at 1200 rpm at 4^0^C. Supernatant was carefully removed, cells were fixed and washed twice in 3:1 methanol:acetic acid (v/v) at 4^0^C. Fixed cells were dropped on clean, humid glass slides and allowed to air dry. For immunoFISH protocols, we followed previously reported methodologies with a few modifications as described. Colcemid treated RPE-1 cells isolated by mitotic shake-off were treated with hypotonic buffer at 3 × 10^5^ cells/ml for 15 min at 37^0^C and ∼3 × 10^4^ cells were spun onto slides at 366 g for 3 min using Thermo scientific Shandon Cytospin 4.

### Fluorescence in situ hybridization

#### Probe denaturation

Centromere enumeration probes were purchased from Cytocell. Hybridization buffer was from Empire Genomics. Hybridization mix was prepared by diluting 1 μL of probe in 5 μL of hybridization buffer and denaturing the mix at 80^0^C and then chilling on ice for 2 minutes. FISH probes used were from Cytocell – LPE001R/G, LPE006R/G, LPE007R/G, LPE011R/G, LPE012R/G, LPE 013R/G-A, LPE018R/G. Labeled BAC probe (RP11-846C20) from Empire Genomics was used as pericentromeric marker for chromosome 21. Whole chromosome paint for Chromosome 10 was from Applied Spectral Imaging.

#### FISH on isolated chromosome spreads

Chromosomes spreads on slides were treated with 100 μg/ml RNase A in 2X SSC (Qiagen, Cat No 19101) for 30 min at 37^0^C and washed in 2X SSC. Slides were dehydrated in an ethanol series of 70%, 90% and 100% and left to air dry. Slides were then denatured at 78^0^C for 2 min in 70% formamide/2X SSC, covered with the pre-denatured hybridization mix and incubated in a humid chamber at 37^0^C overnight.

#### ImmunoFISH on unfixed chromosome spreads

As described in Figure 3, unfixed chromosome spreads were blocked in 10% BSA in KCM buffer (120 mM KCl, 20 mM NaCl, 10 mM Tris-HCl, ph8, 0.5 mM EDTA, 0.1% (v/v) Triton X-100) for 10 min. Slides were incubated in primary antibodies diluted in blocking buffer at 4^0^C for 1 hour and washed in 1X TBST/0.1% Triton X-100 (1X TBST). Primary antibodies were used as follows: mouse anti-CENP-A (MBL, D115-3, dilution 1:100), rabbit anti-CENP-B (Abcam, ab25734, dilution 1:100), rabbit anti-CASC5/KNL1(Abcam, ab222055, dilution 1:100), rabbit anti-HEC1/NDC80 (Thermo Fisher Scientific, PA5-78319, dilution 1:100). Samples were incubated with secondary antibodies at 4^0^C for 1 hour. Secondary antibodies used as follows – Goat anti-Mouse IgG Alexa Fluor™ 568 (Thermo Fisher Scientific LLC, A-11004) and Goat anti-Rabbit IgG Alexa Fluor™ 647 (Thermo Fisher Scientific LLC, A-21245). Slides were post-fixed in 4% paraformaldehyde followed by ethanol dehydration. Slides were denatured at 68^0^C for 3 min with the pre-denatured hybridization mix and incubated in a humid chamber at 37^0^C overnight.

#### ImmunoFISH on fixed chromosome spreads

As described in Figure 3, chromosome spreads were fixed with 1% paraformaldehyde (PFA) in PBS-0.2% Triton-X-100 for 10 mins. Slides were incubated in 50 mM NH_4_Cl for 30 mins and blocked in 3% BSA. Slides were incubated in primary antibodies diluted in blocking buffer at 4^0^C overnight. Primary antibodies used – mouse anti-CENPA (MBL, D115-3, dilution 1:100), rabbit anti-CENP-A (proSci, 30-143, dilution 1:100), rabbit Anti-INCENP (Cell Signalling, 2807S, dilution 1:100), mouse anti-Aurora B (BD Biosciences, 611082, dilution 1:100), rabbit anti topo 2A (Cell Signaling, D10G9, dilution1:100), mouse anti-CENP-E (Abcam, ab5093, dilution 1:100). Secondary antibody treatments as earlier followed by post fixation and ethanol dehydration as earlier.

#### FISH on paraformaldehyde fixed cells

RPE-1 and HCT 116 growing on 18 x 18 mm glass coverslips No. 1.5 (VWR, Cat. No. 48366-205) were fixed in 4% PFA. Cells were permeabilized in 0.5% Triton X-100/PBS and incubated in 20% glycerol/PBS for one hour. Coverslips were flash frozen in liquid nitrogen and allowed to thaw before returning to 20% glycerol. Three freeze-thaw cycles were performed. Cells were washed in PBS and incubated in 0.1 N HCl for 5 min, washed in PBS and incubated in 50% deionized formamide/2X SSC overnight or stored in 50% deionized formamide/2X SSC for up to 3 weeks before hybridization. During hybridization, a drop of pre-denatured hybridization mix was applied to a slide, coverslips containing the fixed cells were inverted onto the hybridization mix, sealed with Cytobond (SciGene, Cat. No. 2020-00-1) and denatured on a heating block set at 80^0^C for 2 minutes. Slides were incubated in a humid chamber at 37^0^C overnight.

#### ImmunoFISH on metaphase-arrested cells

As described in Figures 2, 6, RPE-1 cells were fixed in 4% paraformaldehyde in PTEMF buffer (20 mM PIPES pH 6.6, 10 mM EGTA, 1mM MgCl2, 0.25% Triton X-100). For PICH staining cells were fixed in 4% PFA in PBS and permeabilized in 0.5% Triton X-100. Cells were blocked in 5% BSA in PBST for 30mins. Primary antibodies used were: custom anti-PICH antibody (1:100)^58^, rabbit anti-CENP-B (Abcam, ab25734, 1:100), anti-Aurora A (CST, 14475S, 1:100). Secondary antibodies used as previously. Cells were post fixed in 2% PFA, re-permeabilized in 0.5% Triton X-100, and all steps for FISH fixation of cells were followed, as previously stated. FISH was performed as previously described.

#### Wash stringency

Hybridized chromosome spreads on slides and cells fixed on coverslips were washed identically. The first wash was in 2X SSC at room temperature, followed by three washes in 50% formamide/2X SSC at 45^0^C, followed by three washes in 0.1X SSC at 60^0^C. Samples were rinsed in 4X SSC/0.1% Tween 20 with one final rinse in 2X SSC before mounting in Antifade Mounting Medium with DAPI (Vectashield, Cat. No. H-1200).

### Confocal Microscopy

Chromosomes spreads were imaged on Zeiss LSM 780 and LSM 980 confocal microscopes equipped with GaSaP-PMTs with 405-nm, 488-nm, 561-nm laser lines using a 63x Plan-Apochromat 1.4-numerical aperture oil immersion objective at 2.5x digital zoom. Scanning was performed sequentially (x-y, 512 pixels by 512 pixels [1 pixel ∼ 0.105 μm]), and z-stacks were collected at a step size of 0.34 μm and a pinhole size of ∼0.7 μm (1 AU). Cells were imaged with the Nikon TiE microscope equipped with PlanApo 63x oil immersion objective NA 1.4, Yokogawa CSU-W1 spinning disk, Flash 4.0 sCMOS camera (Hamamatsu), and NIS Elements software. Z-stacks were collected at a step size of 0.34 μm. High content imaging was performed with a Nikon Eclipse Ti2 microscope equipped with a 10-position emission filter wheel, 405, 488 and 545 laser lines, a Yokogawa CSU-W1 spinning disk confocal scanner and a Hamamatsu Orca-Fusion BT sCMOS camera, controlled via NIS-Elements software. Images were acquired through a Nikon Plan-Apochromat 40× water immersion objective (NA 1.2) with exposure times ranging from 50 to 100 ms. The lateral (x,y) pixel size was 0.103 µm, and Z-stacks were captured using a step size of 0.34 µm.

### Image analysis

#### Centromere FISH signal quantification from chromosome spreads

Images were analyzed using several custom open-source FIJI analysis packages available at http://research.stowers.org/imagejplugins/. Source codes can be found at the Github repository https://github.com/jayunruh/Jay_Plugins/blob/master/track_max_not_mask_fast_jru_v1.java. Briefly, for determining the intensities of centromere FISH signals for chromosome spreads, Z-stacks were sum projected and closely cropped to contain chromosomes only from a single metaphase spread. A circular background region 30 pixels in diameter was found on the image and its intensity was subtracted from the whole image. Images were thresholded separately for independent channels and integrated density values were derived. Sometimes more than two signals were obtained from potentially aneuploid spreads; such images were excluded from the analysis.

#### Centromere FISH signal quantification from G1 and G2 arrested cells

Centromere signals from a population of synchronized cells were assessed from sum intensity projection images using FIJI using custom macros available at https://github.com/SidShivanandan/2025_Shivanandan/tree/main. A median filter of 2 pixels was applied to the DAPI channel after which DAPI signals were thresholded manually to define nuclear masks containing a pair of centromere signals. Within the nuclear mask, the centromere arrays for each chromosome pair were analyzed. Centromere FISH signals were manually thresholded to generate centromere masks. Centromere and nuclear masks were associated to determine total intensities of a pair of cells. Nuclei with fewer or more than two centromere masks were omitted from analysis. Data was processed using a custom R script to define dim (lower intensity) and bright (higher intensity) centromeric arrays within every nucleus.

#### Assessment of 3D positions of centromeres

For quantifying the centromere to nuclear center distances in Fig 2C: centromeres were found using a combination of gaussian_laplace and peak_local_max from scipy and scikit-image respectively. The nuclei were segmented with smoothing and otsu thresholding and their centroids identified. Only nuclei in which two centromeres were located were kept, and centromere intensities were quantified after background subtraction and smoothing, with the haplotypes separated by which was brightest. For metaphase alignment in Fig 2D: manually annotated foci for the Aurora A were used as anchors and images rotated and shifted such that their midpoint became origin of the new coordinate system. Bright and dim foci for CEN 1, CEN 7, CEN 6 and CEN 11 were similarly annotated and their foci relative to the origin determined. The distance from each centromere back to its corresponding nuclear center was then calculated. Code available at https://github.com/jouyun/2025_SenGupta_Shivanandan.

#### Quantification of inter-centromere distances from metaphase-arrested cells

Inter-sister distances at centromeres were assessed from maximum intensity projection images using FIJI. A line region of interest (ROI) wide enough to fit the centromere signals was drawn over signals from each homolog. Cohered sisters appeared as a single stretched signal while uncohered sisters appeared as two discrete foci. Line profiles of the intensities were generated and cohered signals were fit to a single gaussian curve while uncohered signals were fit to a double gaussian curve using custom FIJI macros. Source code can be found at – https://github.com/jayunruh/Jay_Plugins/blob/master/fit_traj_single_double_gaus_jru_v1.java.

Single gaussian fits yielded one amplitude and one SD value, double gaussian fits yielded two amplitudes (amp1 and amp2) and two SD (sd1 and sd2) values. x1 was the position of the first peak and x2 was the position of the second peak of intensity. From these parameters, the following values were determined:

Inter-sister distance at centromeres (d) = x2-x1

#### Annotation of dim and bright centromeric arrays in cells

Bright and dim arrays in RPE-1 cells were assessed and annotated using a custom Python script (https://github.com/jouyun/2025_Gupta_Shivanandan/QuantifyAnnotatedLines.ipynb). The script was first validated by visual inspection in Napari using a test set of images to ensure correct annotation and then run unsupervised. Briefly, background was subtracted from images using a tophat filter. Next, a gaussian filter was applied with radius of 2, images thresholded by Otsu thresholding and aggregated segments split by watershed. Intensities of all objects lying on a single ROI were summed. Since each ROI corresponded to a single centromeric array, the summed intensity reflected the total intensity of a single centromeric array out of a pair. The bright array in a cell had higher intensity value compared to the dim array.

Intensities of homologous centromeres in HCT 116 cells were determined by analysis of the area under gaussian fits using the following formulae:

Total intensity = 2 x (amp1)/2 x (sd2)/2 x (√(2 x π)), from single Gaussian fits

or, total intensity = 2 x [(amp1+amp2)/2] x [(sd1+sd2)/2)] x (√(2 x π)), from double Gaussian fits.

Total intensity values were determined for each centromeric array in a cell and used to annotate the bright and dim arrays for HCT 116 cells. The bright array had higher total intensity value compared to the dim array.

#### Quantification of centromere proteins from immunoFISH images

For Figure 3, FISH labeled chromosomes were detected using an Otsu threshold on background subtracted, sum projected 2D images. Background subtraction was performed using tophat filtering from scikit-image. The two brightest objects, as determined by integrated signal intensity, were paired, and the closest CENP-A pair for each was found. CENP-A foci were localized in 3D using a Laplacian of Gaussian filter together with peak_local_max from scikit-image. CENP-A peaks were paired based on shortest distance up to a max distance of 6 pixels. For each of the CENP-A pairs associated with a FISH signal (bright and dim), a line was drawn between the CENP-A foci and all channel intensities in the background subtracted, sum projected 2D image within 6 pixels of this line were quantified. Source codes can be found here – https://github.com/jouyun/2025_SenGupta_Shivanandan

### Statistical analysis and graphs

Statistical analysis was performed using GraphPad Prism v 10.0. Statistical tests are mentioned in figure legends. Graphs were generated using GraphPad Prism v 10.0 or R version 4.0.2.

## Acknowledgements

We thank members of the Gerton lab for discussion and feedback on this work. We thank Leonardo Gomes de Lima for help with centromere repeat analysis. HCT 116 cells were kindly provided by Jan Michel-Peters (IMP, Vienna). PICH antibody was generously provided by Yoshiaki Azuma (KU Lawrence). This research was supported by the Stowers Institute for Medical Research and NIH-NCI under award number R01CA266339 (to JL Gerton).

## Author contributions

A.S.G., S.S. and J.L.G. conceived the project. A.S.G., S.S. and J.L.G. wrote the manuscript with input from all other authors. A.S.G. performed experiments, imaged and analyzed RPE-1 chromosome FISH assays, RPE-1 cohesion experiments. S.S. performed and analyzed cell cycle assays, chromosome missegregation assays and immunoFISH experiments. M.M. performed all experiments, imaging and analysis in HAP1 and HCT 116 cells. A.P.J. performed immunoFISH experiments, imaging and analysis for kinetochore proteins under the supervision of A.S.G. and S.S. J.R.U. and S.A.M. wrote custom ImageJ and Python scripts for analysis of centromere FISH and centromeric protein signals. S.G. provided early access to RPE-1 T2T assembly data. J.L.G. acquired funding for the project.

---

**Figure S1.**
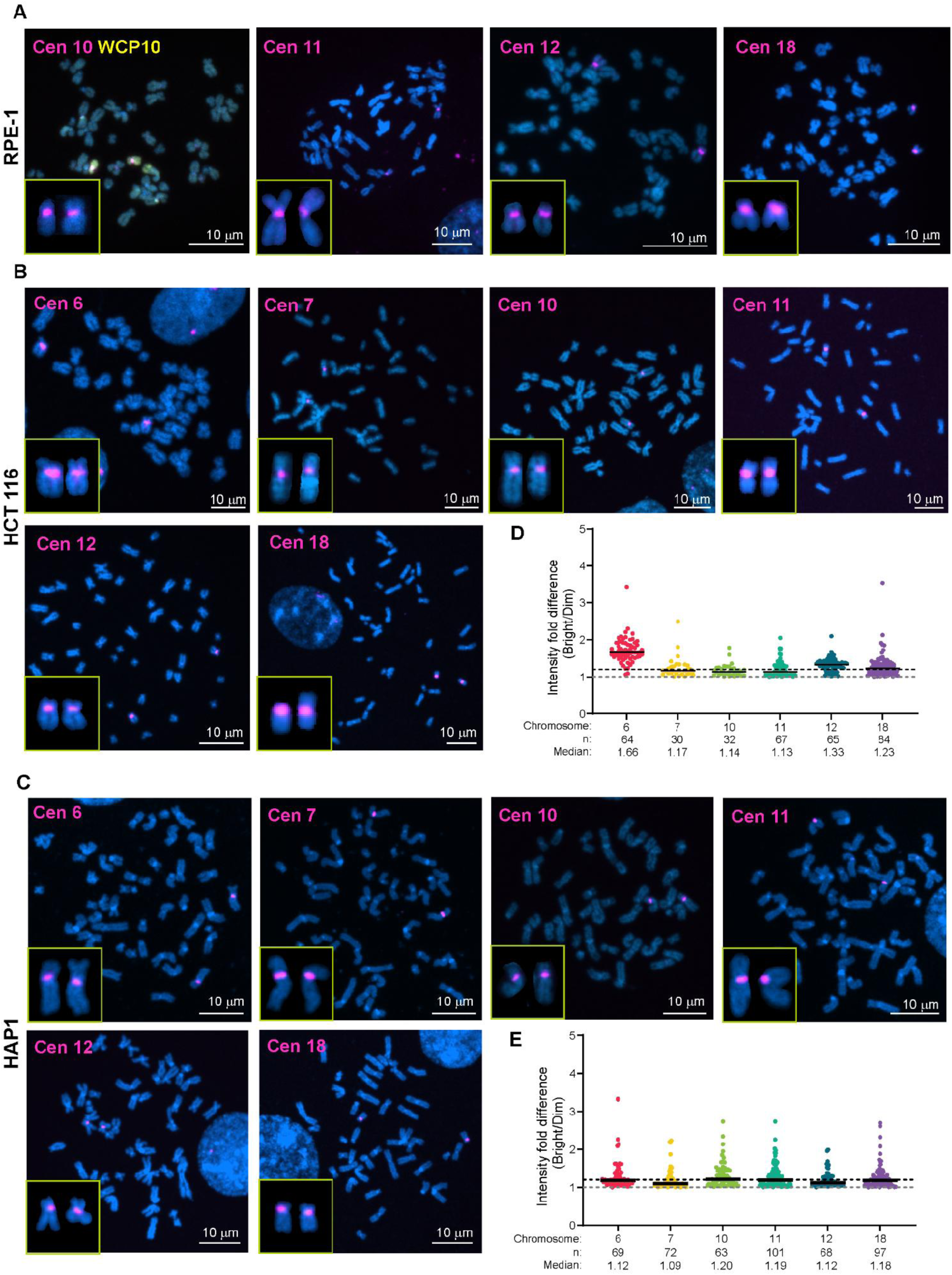
Detection of centromeric array size in a panel of human centromeres in RPE-1, HCT 116 and HAP1 cells. **A-B.** FISH was performed on chromosome spreads from RPE-1, HCT 116 and HAP1 cells. Representative images of all centromeres that were assessed by FISH in each cell line are shown. Insets show a pair of homologous chromosomes from every spread. **C- D.** Fold difference in centromere FISH signal intensity from homologous chromosome pairs in (C) HCT 116 and (D) HAP1 cells. Bar denotes the median. Fold-change of 1 denotes no difference in centromeric array size between homologs. Chromosomes with median higher than 1.2-fold (dotted line) indicate homologs with detectable variation in centromeric array size.

**Figure S2.**
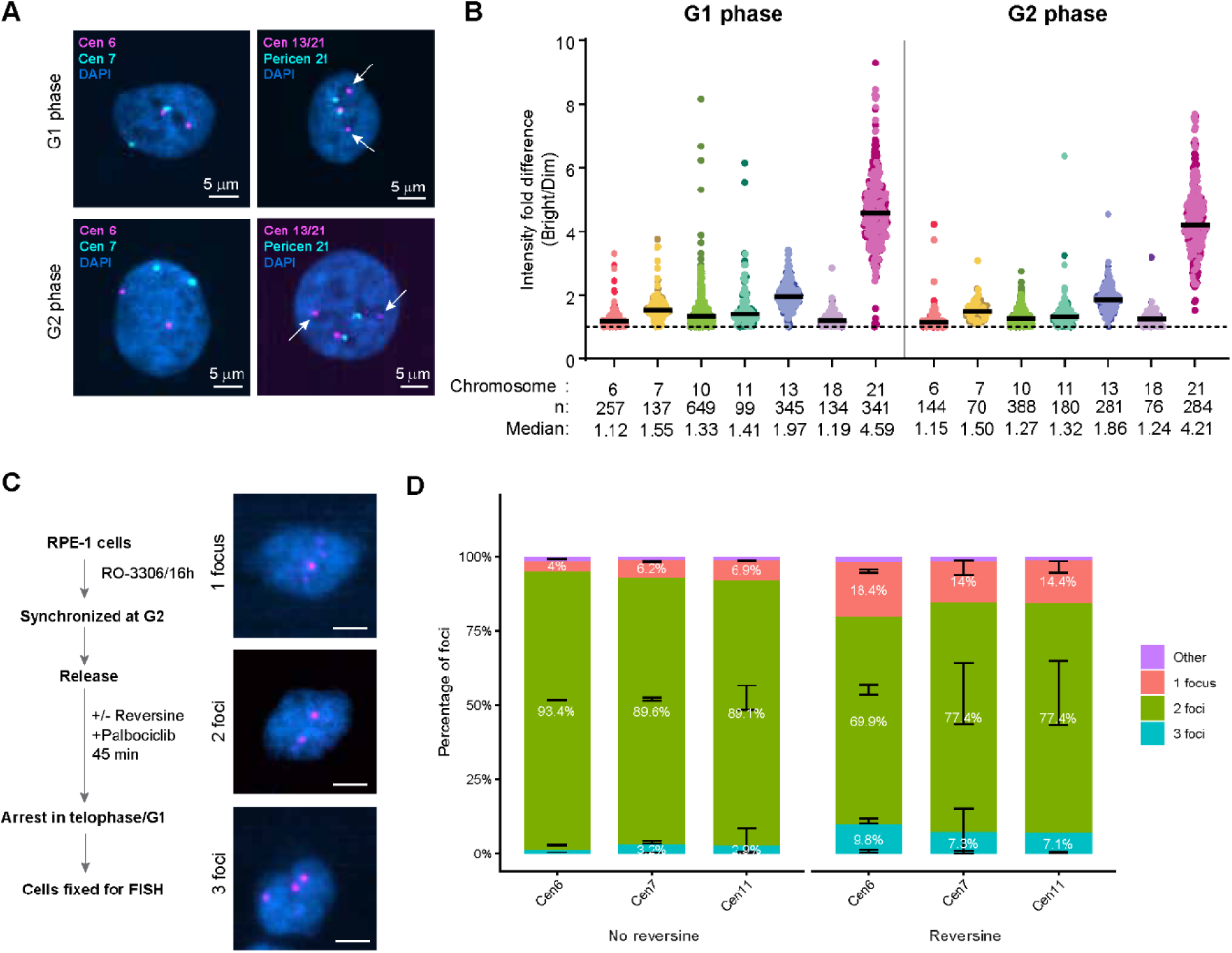
Chromosome segregation analysis in RPE-1 cells. **A.** RPE1 cells were synchronized in G1 or G2 phase and centromeres were assessed by FISH. Representative images of centromere FISH signals for a few selected chromosomes 6, 7, 13 and 21 are shown. Chromosome 21 is distinguished by a chr21-specific pericentromeric probe. **B.** Fold difference in centromere FISH signal intensity from homologous chromosome pairs in G1 and G2 arrested RPE-1 cells. Bar denotes the median. **C.** RPE-1 cells were synchronized in G2/M and released in medium with or without the spindle checkpoint inhibitor reversine and then arrested before G1 to allow a single chromosome segregation event. Centromere FISH signals in RPE-1 cells were assessed by high content imaging analysis. Cells revealed one, two, or three centromere FISH signals. Cells with two centromere signals indicate an accurate chromosome segregation event whereas a deviation from two signals indicates missegregation. Scale bars denote 5 μm. **D.** Distribution of RPE-1 cells showing 1, 2, 3 or other number of centromere signals after one round of chromosome segregation. Error bars represent standard deviations from the mean. The percentage of cells with three foci in cells with an intact spindle checkpoint is low (<5%) for all assessed chromosomes – 6, 7, and 11. Upon reversine treatment, all assessed chromosomes show more than 5% of cells with three foci, allowing the assessment of size-based bias in centromere performance during chromosome segregation.

**Figure S3.**
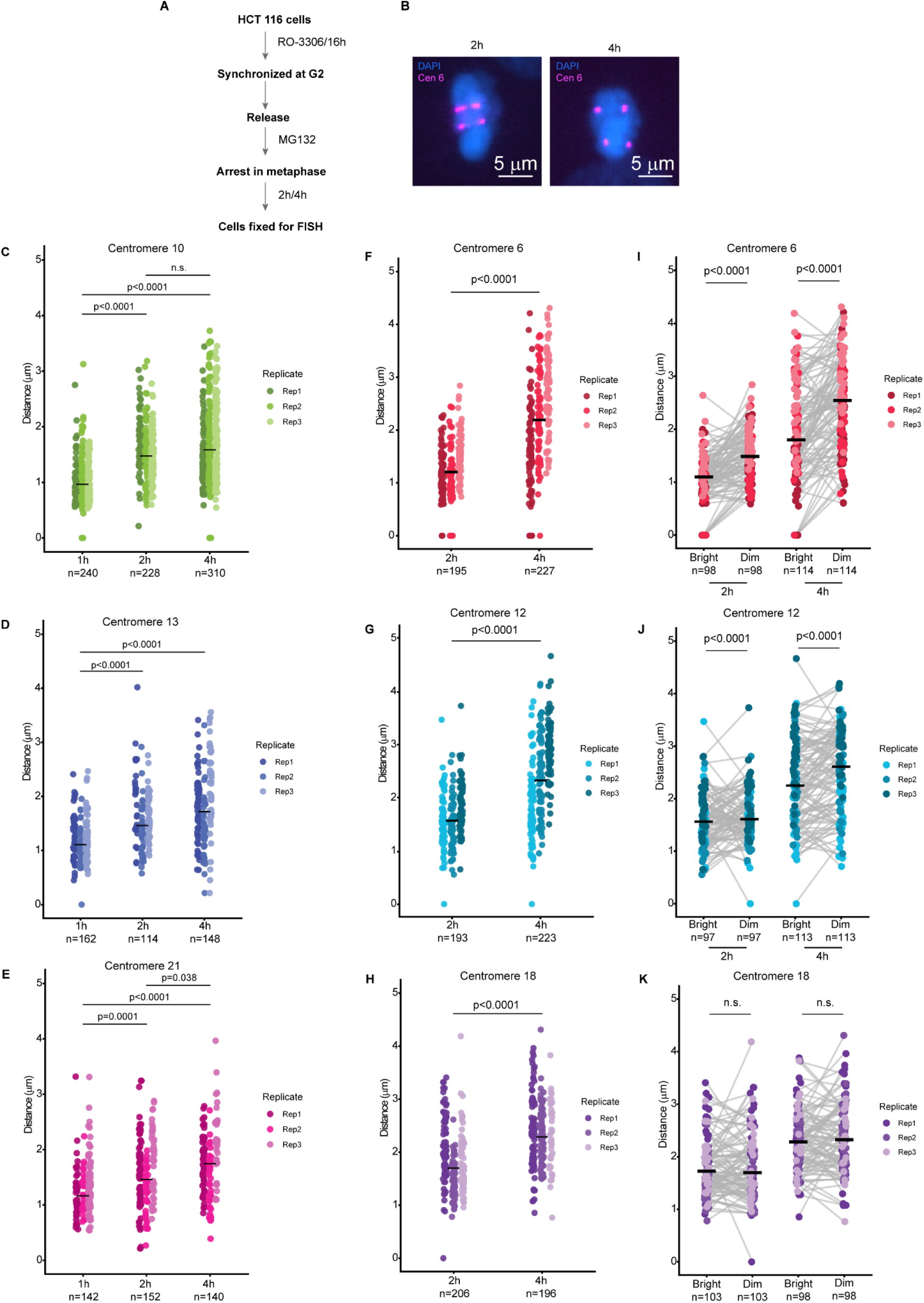
Centromere array size-based bias in cohesion upon prolonged metaphase arrest in RPE-1 and HCT 116 cells. **A.** Similar to RPE-1 cells shown previously, HCT 116 cells were synchronized in G2/M and released in medium containing the proteasome inhibitor MG132 to arrest cells in metaphase for 2h or 4h and then fixed for centromere FISH. **B.** Representative images of HCT 116 centromeres showing cohesion fatigue at both 2h (left panel) and 4h (right panel) of metaphase arrest. **C-E.** RPE-1 cells were arrested in metaphase after release from synchrony in G2 phase. Increasing inter-centromere distances for centromere 10, 13 and 21 at 1h, 2h or 4h of metaphase arrest showing loss of cohesion as a function of time. Bars indicate median distance. Statistical significance tested by Kruskal-Wallis test. **F-H.** HCT 116 cells were arrested in metaphase after release from synchrony in G2/M phase. Increasing inter-centromere distances for centromere 6, 12 and 18 at 2h or 4h of metaphase arrest showing loss of cohesion as a function of time. Bars indicate median distance. Statistical significance tested by Kruskal-Wallis test. **I-K.** Pairwise comparison of inter-centromere distances at bright and dim centromeres at 2h and 4h of metaphase arrest from chromosomes 6, 12, and 18 in HCT 116 cells. Dim centromeres of chromosomes 6 and 12 show higher inter-centromere distances in HCT 116 cells. Centromere 18 homologs with undetectable size variation do not show this cohesion bias. Bars indicate median distance. Statistical test used is Wilcoxon test. n denotes the number of cells assessed in every condition from three biological replicates. n.s. denotes not significant.

**Figure S4.**
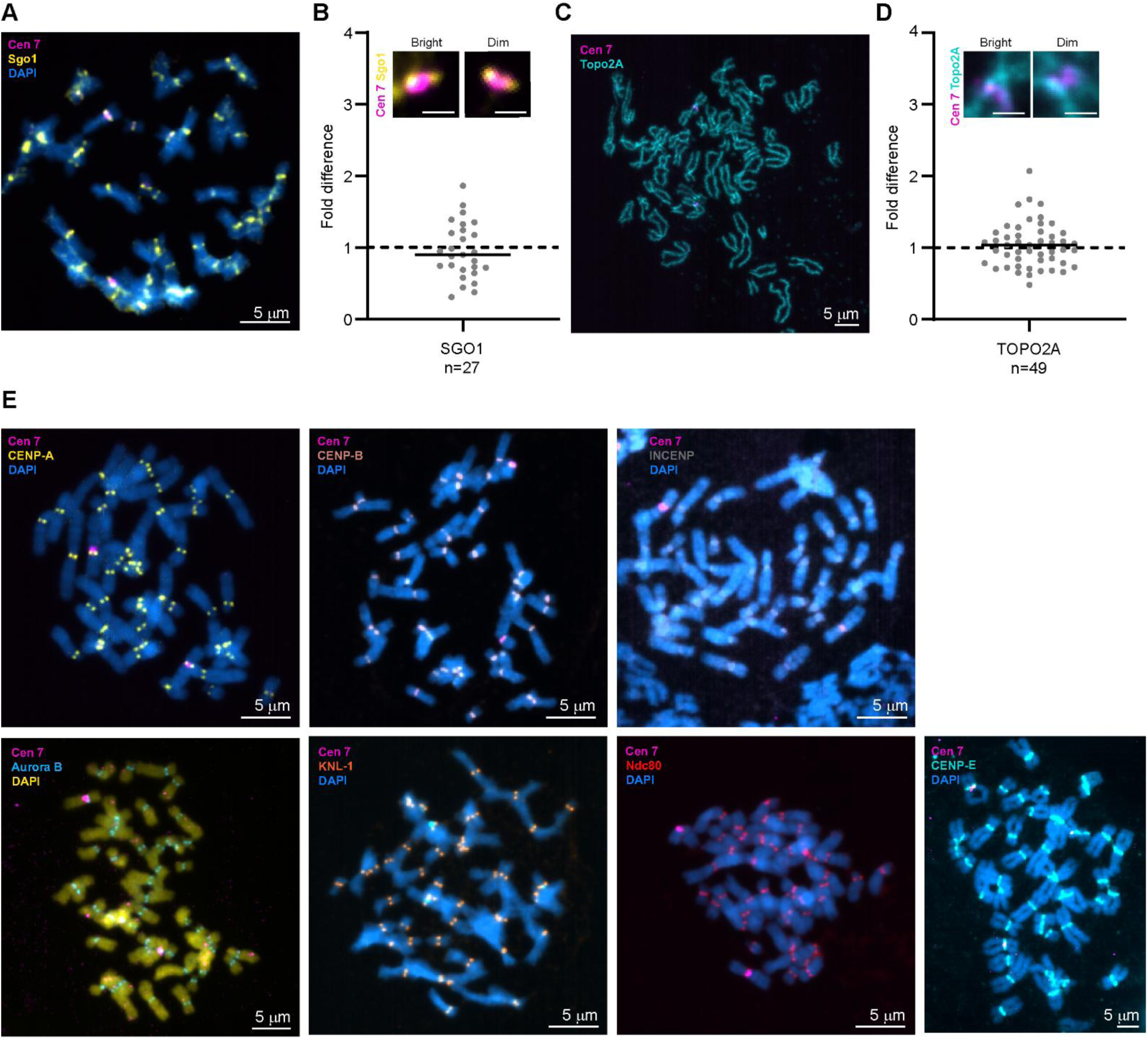
Enrichment of non-kinetochore and kinetochore proteins on RPE-1 chromosome spreads. **A.** ImmunoFISH for SGO1 and centromere 7 in RPE-1 chromosome spreads. **B.** Fold-difference of SGO1 intensities on bright and dim arrays of centromere 7. No significant difference was observed. **C.** ImmunoFISH for TOPO2A and centromere 7 in RPE-1 chromosome spreads. Inset scale bars denote 1μm. **D.** Fold-difference of TOPO2A intensities on bright and dim arrays of centromere 7. No significant difference was observed. **E.** Representative images of immunoFISH for all kinetochore components assessed in this study. Because Aurora B is consistently color-coded blue, we used yellow for DAPI in the image with Aurora B.

**Figure S5.**
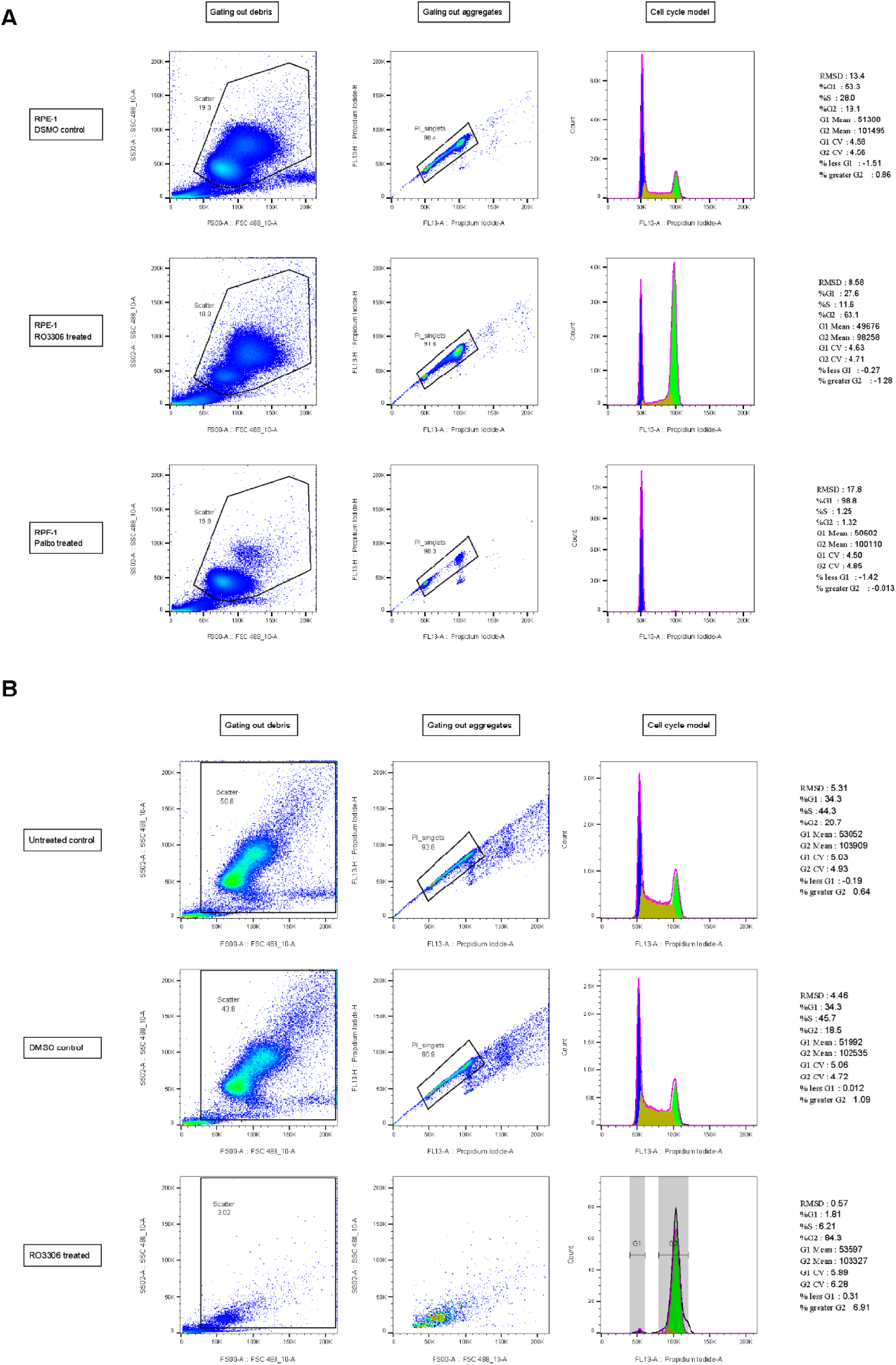
Cell cycle analysis after synchronization. **A.** Cell cycle profiles are shown for RPE-1 cells treated with RO-3306 or Palbociclib to synchronize cells in G2/M or G1 phases, respectively. After treatment, cells were fixed and stained with propidium iodide for cell cycle analysis by flow cytometry. **B.** Cell cycle profiles of HCT 116 cells are shown, treated with RO-3306 to arrest in G2/M phase, DMSO, or untreated.

**Figure S6.**
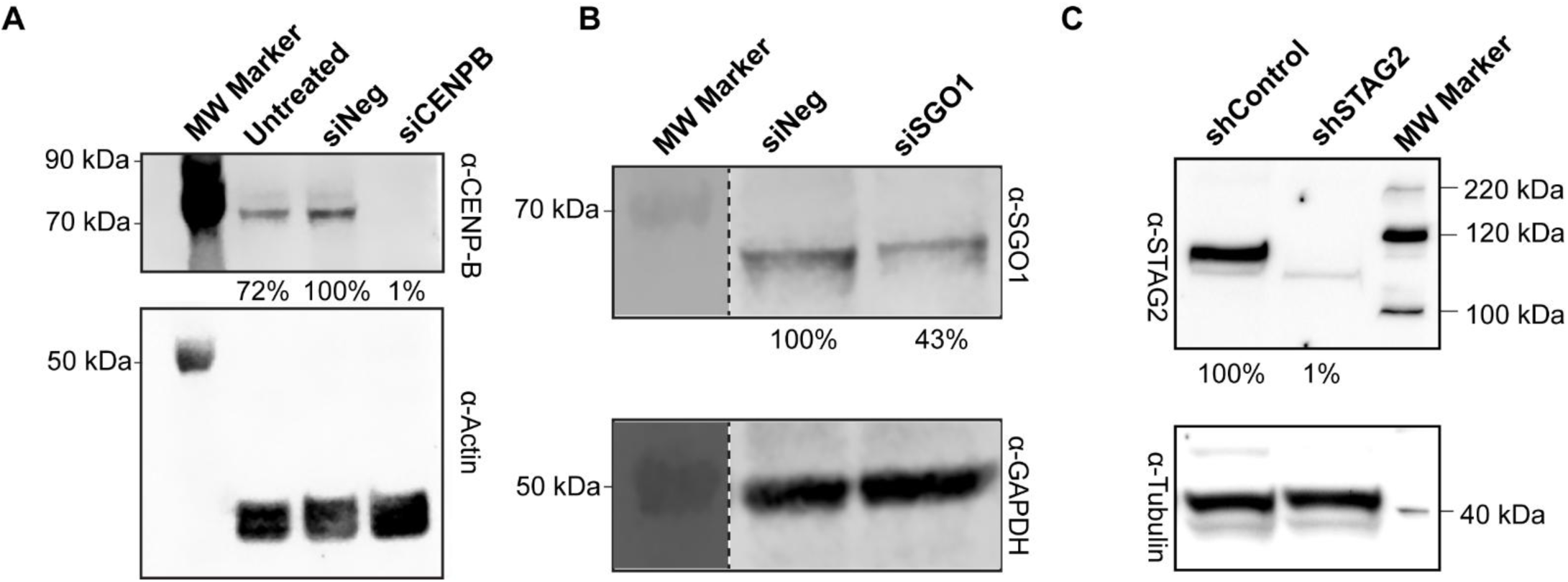
Assessment of knockdown efficiencies of centromeric proteins in RPE-1 cells. **A-C.** Western blots showing the knockdown efficiency of **(A)** CENP-B after 48 hours, **(B)** SGO1 after 24 hours and **(C)** STAG2 upon constitutive knockdown in RPE-1 cells. Actin, GAPDH and β-tubulin served as loading controls. Equal amounts of total protein were loaded per lane. Expression levels of CENP-B, SGO1 and STAG2 are shown below the blot normalized to respective loading controls. The lane containing molecular weight markers from the corresponding blot (B) was spliced to the lanes shown, demarcated by dotted lines.

